# Structural Insights into the ATP-dependent Activation of NOD-like Receptor with Pyrin 3 (NLRP3) Protein by Molecular Dynamics Simulation

**DOI:** 10.1101/2023.05.03.539258

**Authors:** Christina F. Sandall, Justin A. MacDonald

## Abstract

The inflammasome-forming NOD-like receptor containing pyrin-3 (NLRP3) protein is a critical player in the innate immune responses to cellular danger signals. New structural data of NLRP3 provide a framework to probe the conformational impact of nucleotide binding. In this study, microsecond molecular dynamics (MD) simulations were used to detail information on the unique structural conformations adopted by NLRP3 with ATP or ADP binding. Sampling convergence reflected a high degree of confidence in the MD simulations as shown by RMSD and protein-nucleotide concordance, favourable overall MM-PBSA ligand binding energies for both nucleotides and low cosine coefficients of the principal eigenvectors obtained with essential dynamics (ED) analysis. NLRP3-ADP simulations provide relatively stable conformations with few global rearrangements as shown by decreased protein RMSD, Rg, SASA, and solvent accessibility for the ADP-bound structure. In contrast, ATP binding induced increased flexibility and resulted in substantive conformational changes to the NLRP3 structure. Binding of ATP was thermodynamically favourable as shown by the ΔG_solv_ and MM-PBSA calculations of complex free energies, and these NLRP3-ATP simulations resulted in similar structural transitions as observed in the activated NLRC4 empirical structure. Lastly, the active conformation of NLRP3 critically depends on hinging between the HD2 and LRR domains, whereby ATP binding drives local conformational changes that are conveyed to the global structure.

## 1. INTRODUCTION

Human nucleotide-binding domain and leucine-rich repeat-containing (NLR) family members are characterized by a central nucleotide-binding NACHT domain that displays nucleoside triphosphatase (NTPase) activity (Ting et al. 2008). The NACHT domain derives its name from several of the proteins first identified to contain it (i.e., <u>NA</u>IP (Neuronal Apoptosis Inhibitory Protein), <u>C</u>IITA (class II major histocompatibility complex transactivator), <u>H</u>ET-E (heterokaryon incompatibility protein from *Podospora anserina*) and <u>T</u>EP1 (telomerase protein component 1)) (Koonin and Aravind 2000). Conservation of the NACHT domain in numerous proteins involved in innate immunity reflects the vital functional importance of this structural fold (Danot et al. 2009).

Comparative analyses reveal sequence conservation within the NACHT domain that designates NLRs as members of the <u>S</u>ignal <u>T</u>ransduction <u>A</u>TPases with <u>N</u>umerous <u>D</u>omains (STAND) clade of the AAA+ ATPase superfamily of P-Loop NTPases (i.e., ATPases-associated with various cellular activities) (Koonin and Aravind 2000; Leipe, Koonin, and Aravind 2004; Snider and Houry 2008). Early studies of STAND ATPases, including plant R proteins (Meyers et al. 1999), *Caenorhabditis elegans* cell death gene CED-4 (Chinnaiyan et al. 1997) and the APAF1 apoptosome (Zou et al. 1999) provided the first indication that nucleotide binding by NLRs was likely to be a regulated process integral to the control of inflammasome assembly.

The NLRs with pyrin domain (NLRPs) comprise an important cytosolic sub-group of NLRs that respond to cellular danger molecules. NLRP3 is the most extensively studied NLRP protein and is the prototypical inflammasome-forming member of the family (Xu and Nunez 2022). Activating stimuli for NLRP3 are famously diverse and include K^+^ efflux, Cl^−^ efflux, Ca^2+^ influx, mitochondrial DNA (mtDNA), alum, reactive oxygen species (ROS), uric acid crystals implanted biomaterials or medical devices, asbestos, silica, and calcium oxalate as well as a variety of metabolic constituents including cholesterol crystals and succinate accumulation (Banerjee et al. 2022; Holley and Schroder 2020; Olona, Leishman, and Anand 2022; Shirasuna, Karasawa, and Takahashi 2019). Historically, two distinct and seemingly compulsory steps were characterised for assembly of the NLRP3 inflammasome: an initial transcriptional priming step and a second ATP-dependent oligomerization and assembly step (Platnich and Muruve 2019; Kelley et al. 2019; McKee and Coll 2020). Once assembled, the NLRP3 inflammasome elicits auto-proteolysis and activation of pro-caspase-1, which in turn triggers the maturation and secretion of pro-inflammatory mediators to the surrounding extracellular milieu (Broz and Dixit 2016; Weber et al. 2020).

It is generally accepted that ATP hydrolysis is the fundamental enzymatic property of NLRP3 that enables the protein to initiate oligomeric inflammasome assembly and downstream effector activation. Examining the primary sequence of the NACHT domain of NLRP3 not only reveal multiple conserved regions for ATP-binding (e.g., Walker A, Walker B and PhhCW motifs as well as the winged-helix domain, WHD) but also several distinct amino acids that may offer a functional selectivity (Maharana, Panda, and De 2018; MacDonald et al. 2013; Proell et al. 2008). For example, Ser295 in human NLRP3 can be phosphorylated to attenuate inflammasome activity (Sandall and MacDonald 2019). Moreover, mutations within the ATP-binding region of NLRP3 that abrogate ATPase activity suppress the maturation of IL-1β (Duncan et al. 2007). Bioluminescence resonance transfer (BRET) experiments have also revealed conformational changes in NLRP3 that occur upon activation with the promotion of an ‘open’ or ‘active’ state of the NACHT domain (Tapia-Abellan et al. 2019).

Newly deposited structural data for NLRP3 now provide a framework to probe the conformational effects of nucleotide binding. In this study, we investigated whether ATP ligand occupancy could instigate activating conformational changes in the monomeric NLRP3 protein. Multiple molecular dynamics (MD) simulations of both ADP-bound and ATP-bound NLRP3 were computed and analysed to reveal an atomic-level picture of protein motion and a representation of the free-energy landscape (FEL) of protein conformations. The results reveal ATP binding with NLRP3 to be thermodynamically favourable, to induce increased flexibility in the NLRP3 backbone and to drive distinct conformational changes in protein topology as compared with NLRP3-ADP. Moreover, ATP-bound NLRP3 underwent comparable structural transitions to activated NLRC4 as a sole result of bound ATP.

## 2. METHODS

### 2.1 Molecular Dynamics: NLRP3 Protein System Preparation

*Software* - Molecular dynamics were run with GROMACS (v2020.4; Abraham et al. 2015), and FoldX was obtained with an academic license (Schymkowitz et al. 2005). Structural figures were generated with the Pymol Molecular Graphics System (Pymol v2.0), and analysed with the UCSF Chimera Visualization System, Visual Molecular Dynamics (VMD), Avogadro Molecular Editor and the Swiss-PdbViewer (DeepView). Statistical analyses were run in R (v4.0.2), PRISM GraphPad (v9.3), or with Real-Statistics (v7.6). Plots and illustrations were generated using GraphPad, Microsoft PowerPoint and Adobe Photoshop.

*Homology Modelling* - The starting structure of human ADP-bound NLRP3 in complex with NEK7 was obtained from the Protein Data Bank (PDB ID: 6NPY; Sharif et al. 2019), and the coordinates of the NEK7 chain were removed. The structural coordinates of NLRP3 include amino acid residues 135-1036, with the N-terminal PYD domain (residues 1-93) deleted. Due to poor density capture, the 3.8 Å resolution cryo-EM deposition of NLRP3 contains seven gaps in the structural data for residues D153-V162 (gap 1), S179-P202 (gap 2), G456-H460 (gap 3), E538-T544 (gap 4), D557-L562 (gap 5), S590-K615 (gap 6), and N656-L681 (gap 7). Initial NLRP3 protein homology models were generated using SWISS-MODEL and MODELLER servers (**Figure 1**) that rely upon the comparative modelling engine ProMod3 (Studer et al. 2021) based on OpenStructure (Biasini et al. 2013), and the MODELLER comparative modelling algorithm based upon the satisfaction of spatial restraints in the CHARMM-22 molecular mechanics forcefield (Vanommeslaeghe et al. 2010) as well as a representative set of high-quality structures from SCOP (Andreeva et al. 2020; Vanommeslaeghe et al. 2010), respectively. An initial quality assessment of 11 models using the built-in assessment scoring functions of SWISSMODEL and MODELLER revealed scores lower than experimental structures on average. To improve the structural accuracy and physical realism of the initial NLRP3 homology models, each was submitted to protein structure refinement servers 3Drefine (Bhattacharya and Cheng 2013) and ReFOLD (Shuid, Kempster, and McGuffin 2017) for integrated quality assessment. Models with scores outside expected parameters were eliminated, and the remaining models were ranked by summated scores. The top 11 models met expected quality criteria and were selected for further scoring and analysis by MolProbity (Williams et al. 2018), ERRAT (Colovos and Yeates 1993), PROVE (Pontius, Richelle, and Wodak 1996), Verify3D (Eisenberg, Luthy, and Bowie 1997) and PROCHECK (Laskowski et al. 1996). Following minimization and re-scoring, three complementary models, which met all evaluation criteria, were selected for preliminary MD simulations (300 ns; performed in quadruplicate) following system preparation, including solvation, energy minimization and equilibration. Ultimately, a single NLRP3ΔPYD protein backbone model was chosen for direct comparison of ADP and ATP ligand occupancy. The selected model yielded the highest validation scores in a composite indicator scoring matrix constructed with model quality assessment data from SAVES (Ginalski et al. 2003; Yang, Roy, and Zhang 2013), MolProbity (Williams et al. 2018) and ModFold (McGuffin et al. 2021) and presented the most stable trajectories over the 300 ns simulation time. This composite model approach of combining complementary pipelines was routinely employed to improve modelling accuracy, fold recognition and ligand binding site predictions (Ginalski et al. 2003; Yang, Roy, and Zhang 2013).

**Figure 1.**
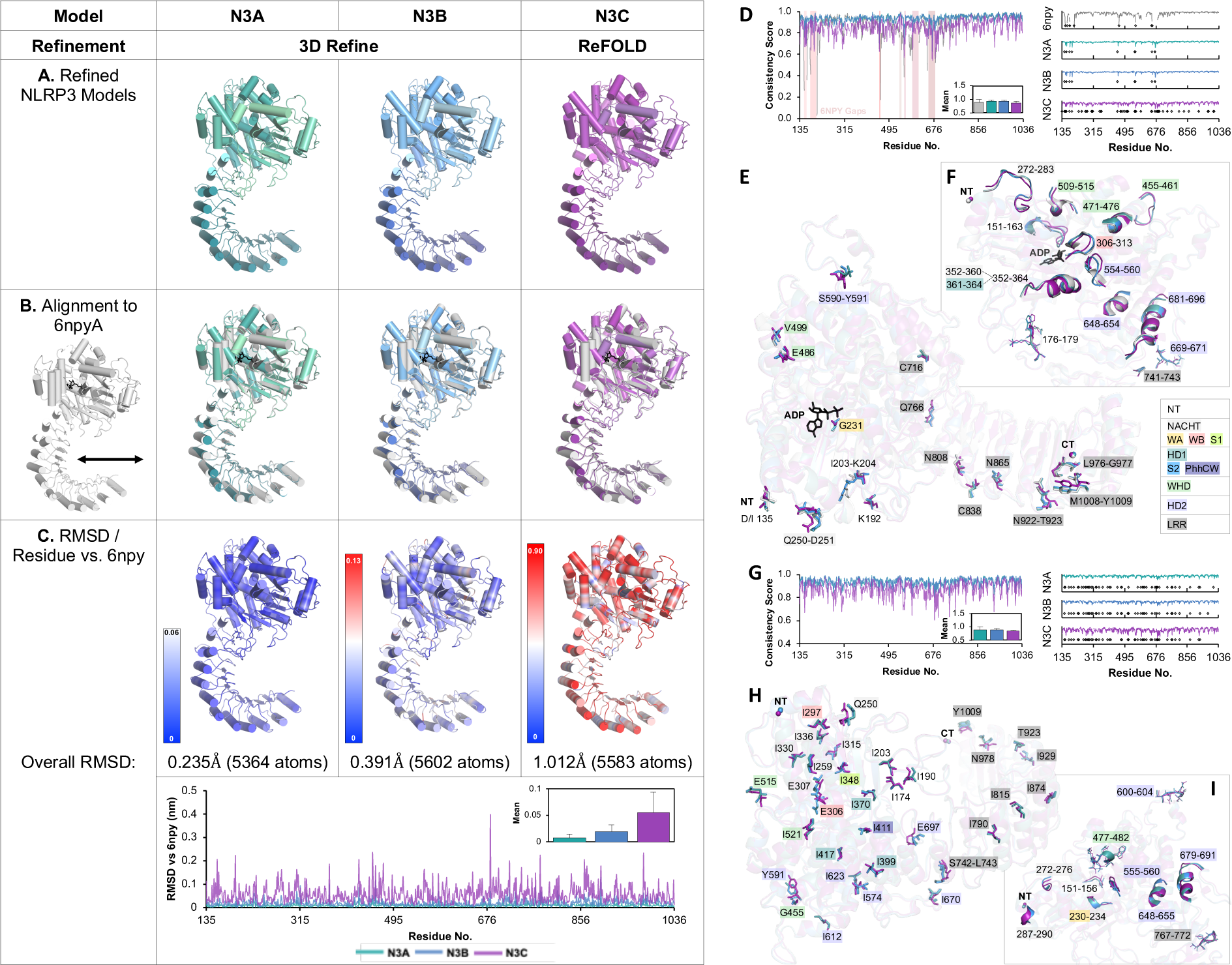
Ensemble Consistency of Homology Models and the Cryo-EM Template Structure of NLRP31′PYD. In (**A-C**), RMSD comparison of three refined NLRP3 homology models with the 6NPYA. Models N3A and N3B were refined with 3D-Refine, and N3C was refined with ReFOLD. (**A**) Cartoon representations of the 3 refined backbone models are provided in (**A**). Each model was aligned with 6NPYA (coloured in gray, **B**) and coloured in a spectrum of blue (0 Å) to red (0.9 Å) by individual residue RMSD, represented graphically to the left of each structure in (**C**). The per residue ensemble consistency scores of NLRP31′PYD homology models as well as 6NPYA are shown in (**D**), with individual data series to the right, and mean scores ± SD included in the inset. Gaps in the 6NPYA structure are represented by gridlines in red. Consistency scores were calculated via the SWISS-MODEL structural comparison system using all other selected structures as reference. Lower scores represent regions of low consistency, and local minima also below 50% of the range of data are represented by diamonds below the individual data sets. Short 1-2 residue stretches (**E**) and regions longer than 2 amino acids (**F**) with consistency scores below 50% of the ranges of data shown in panel D are represented as opaque sticks on transparent cartoon alignments. The gap regions artificially decrease consistency scores, so the ensemble consistency of NLRP31′PYD homology models in the absence of 6NPYA are provided in (**G**). Short 1-2 residue stretches (**H**) and regions longer than 2 amino acids (**I**) with consistency scores below 50% of the ranges of data shown in panel G are represented as opaque sticks on transparent cartoon alignments.

*Nucleotide and Mg^2+^ ion Docking* - To analyse the effect of nucleotide identity on conformational change via the MD simulations, each model was docked with ADP and two conformations of ATP. First, the nucleotide ligand structure from the 6NPY deposition was aligned to each new homology structure using the Pymol alignment algorithm, resulting in RMSD values of 0.235 Å (over 5364 aligned atoms), 0.391 Å (5602 atoms) and 1.012 Å (5583 atoms) for the three models. Next, coordinates for the three *de novo* aligned 6NPY-ADP models were submitted as the templates for ligand alignment with the query molecule ATP (PubChem CID: 5957) using the atom-level small-molecule structural alignment server, LS-align (Hu et al. 2018). Molecular alignment of the ATP ligand was completed with Flexi-LS-align for flexible structure comparison. A Mg^2+^ ion was docked into the final ADP/ATP bound structures with the MIB (Metal Ion-Binding Site Prediction and Docking Server; Lin et al. 2016) and coordinated by the β or γ-phosphate groups of ADP or ATP, respectively, as reported for other AAA+ proteins.

*System Solvation and Neutralization* - The protonation states of the titratable sidechains were determined using PROPKA (v3.1) at a pH of 7.4 (Olsson et al. 2011; Sondergaard et al. 2011). Default tautomeric states were assigned to all residues with the exceptions of Asp804 (protonated, ASH), His492 and His522 (δ-protonated, HID). All remaining His residues were ε-protonated (HIE). Model systems were solvated in a dodecahedron of TIP3P water molecules with 1 nm periodic boundaries. Next, the system was neutralised and brought to a final ionic concentration of 0.15 M. The total number of atoms in the systems was 172,195 for NLRP3ýPYD-ADP and 172,197 for NLRP3ýPYD-ATP. Final systems were developed using CHARMM-GUI (Brooks et al. 2009; Jo et al. 2008).

*System Minimization and Equilibration for MD Production Runs* - MD production runs were performed using GROMACS (v2020.4; Abraham et al. 2015) with the refined all-atom additive CHARMM36m protein force field (Vanommeslaeghe et al. 2010; Huang et al. 2017). System energies were minimised with the steepest descent method until the maximal force in the system was less than 1000 kJ/mol·nm. The system was equilibrated using constant-substance, constant-volume, constant temperature ensemble (NVT) then constant-temperature, constant-pressure ensemble (NPT) for 1000 ps with 1 fs integration time steps. The positions of all non-hydrogen atoms in the protein and the nucleotides were constrained during the equilibration and minimisation runs to stabilise the protein structure and enforce the nucleotide-Mg^2+^ interaction within the nucleotide-binding domain. All bonds to hydrogen atoms were constrained using the LINCS algorithm (Hess et al. 1997). The temperature of the simulations was set at 300 K using the modified Berendsen thermostat with a time constant of 0.1 ps. For the NPT simulations, the pressure was set at 1 bar using isotropic coupling with a time constant of 1.0 ps. The particle-mesh Ewald method (Darden, York, and Pedersen 1993) was used to calculate electrostatic forces with a real space cut-off distance of 1.2 nm, and Lennard-Jones forces were smoothly switched off from 0 to 1.2 nm. Finally, position restraints were released, and the molecular systems were simulated for 1000 ns in two replicates with a time step of 2.0 fs using different seed values and recording the coordinates at every 10 ps.

*MD Trajectory Analyses* - Following the MD production runs, the resultant trajectories were concatenated, periodic boundary conditions were removed, and inbuilt GROMACS functions were used to calculate the RMSD (root mean square deviation; gmx_rms), RMSF (root mean square fluctuation; gmx_rmsf), Rg (radius of gyration; gmx_gyrate), and SASA (solvent-accessible surface area; gmx_sasa). Geometric clustering was calculated using the GROMOS algorithm (Daura et al. 1999). NACHT-domain volumes of the medoid structures of the largest GROMOS clusters were computed with ProteinVolumev1.3 (Chen and Makhatadze 2015). Angle and displacement of the NACHT and LRR domains were calculated in Pymol with a script obtained from the open-source Pymol repository [orientations.py]. Protein-nucleotide hydrogen bond occupancies were calculated with the VMD HBonds plugin (Humphrey, Dalke, and Schulten 1996), where occupancies indicate the total number of hydrogen bonds as a percentage of frames in the concatenated trajectories. The concatenated trajectories were analysed every 10^th^ frame over a total of 20,000 frames. Hydrogen bonds were identified between polar atoms less than 3 Å apart with a 20° angle cut-off. Internal protein-protein hydrogen bonds were analysed with the object-oriented Python library MDAnalysis on the same number of frames (Michaud-Agrawal et al. 2011).

### 2.2 Model Selection

After MD production runs were completed for all models with docked nucleotide ligands, descriptive statistics were assigned for each model to investigate the overall structure of the indicators and assess the suitability of the data sets, ensuring gaussian distributions and a lack of skewness, kurtosis or outliers. Next, the scoring metrics were normalised by standardisation (i.e., computing Z-scores for each metric), and the model Z-scores were calculated on the metric composite to assess each model’s aggregated scores and ranks.

### 2.3 Principal Component Analysis and Free Energy Landscape Calculations

Principal component analysis (PCA) of the trajectories was employed to resolve the largest amplitude of collective motions in the trajectories (Maisuradze, Liwo, and Scheraga 2009). In brief, a diagonalized covariance matrix was constructed from the atomic fluctuations in each trajectory wherein a collection of principal components (i.e., eigenvectors) that describe the collective motion direction in conformational space were obtained with corresponding eigenvalues (i.e., the amplitude of fluctuations along the eigenvector). Essential dynamics (ED) calculations were performed with inbuilt GROMACS functions gmx_covar and gmx_anaeig. Free energy landscape (FEL) calculations were performed using the GROMACS function gmx_sham to calculate the multidimensional free energies along the first two eigenvectors (PC1 and PC2). 2D and 3D FEL plots were generated with modified python scripts [fel3d.py & fel2d.py; www.dlcompbiobiotech.com; Mittal et al. 2021). The ΔG_solv_ was first estimated with the in-built GROMACS function gmx_sasa to compute the solvent-accessible areas (Floris and Tomasi 1989).

The Molecular Mechanics (MM) energies were combined with the Poisson–Boltzmann Surface Area (MM-PBSA) approach to calculate more accurate free energies for binding of the nucleotide ligand (Wang et al. 2019). The gmx_MMPBS algorithm was used to calculate the contributions of various interactions to the overall free energy, including MM changes in the gas phase (ΔE_MM_), additional measures of the free energy of solvation (ΔG_solv_), the enthalpy of ligand binding (ΔH), and conformational entropy (−TΔS) upon ligand binding (Duan, Liu, and Zhang 2016; Valdes-Tresanco et al. 2021). To ensure congruence with the MD simulations during MM-PBSA calculations, the ionic strength (μ) was set to 0.15M, atomic radii for PB calculations were derived from average solvent electrostatic charge distributions within the CHARMM-22 forcefield, the dielectric constant (ε_in_) was set to 3, and the dielectric interface was implemented with a level-set based algebraic method (Wang et al. 2012). Entropic changes (TΔS) were calculated by averaging the interaction entropy (IE) term from 100 snapshots taken at the energy minima for each structure: 173-183ns (ADP) and 106-116 ns (ATP), which was subtracted from the Δ_TOTAL_ term to obtain the absolute binding energy (ΔG_bind_) in kcal mol^−1^.

## 3. RESULTS

### Trajectory Analyses of ADP- and ATP-bound NLRP31′PYD

The NLRP31′PYD model bound with either ADP or ATP was subjected to two MD simulations of 1000 ns each in an explicit aqueous solution; replicas were concatenated and analysed in aggregate. **Figure 2A** shows the time progression of the protein RMSD of both the ADP- and ATP-bound trajectories with respect to the equilibrated starting structure. As expected, both simulations displayed relatively large initial structural deviations as the trajectories converged. However, stable RMSD values were maintained after 36.8 ns and 40.3 ns, plateauing at 24.5 Å and 28.7 Å for the ADP and ATP bound models, respectively. The ADP- and ATP-bound forms deviated from the minimised starting structure by an average of 23.9 Å and 27.4 Å, respectively (**Figure 2A**), indicating that binding of ATP resulted in a more flexible structure. The ATP bound structure underwent a set of more extensive conformational changes as compared with ADP. It is evident that the simulation intervals were sufficient to capture structural transitions, as demonstrated by the RMSD histograms in **Figure 2B**, where distinct peaks suggest dissimilar global conformations of the protein backbone. The ATP-bound trajectory displayed at least two distinct conformations, while the ADP-bound model quickly converged upon a single RMSD value and was relatively stable over the trajectory.

**Figure 2.**
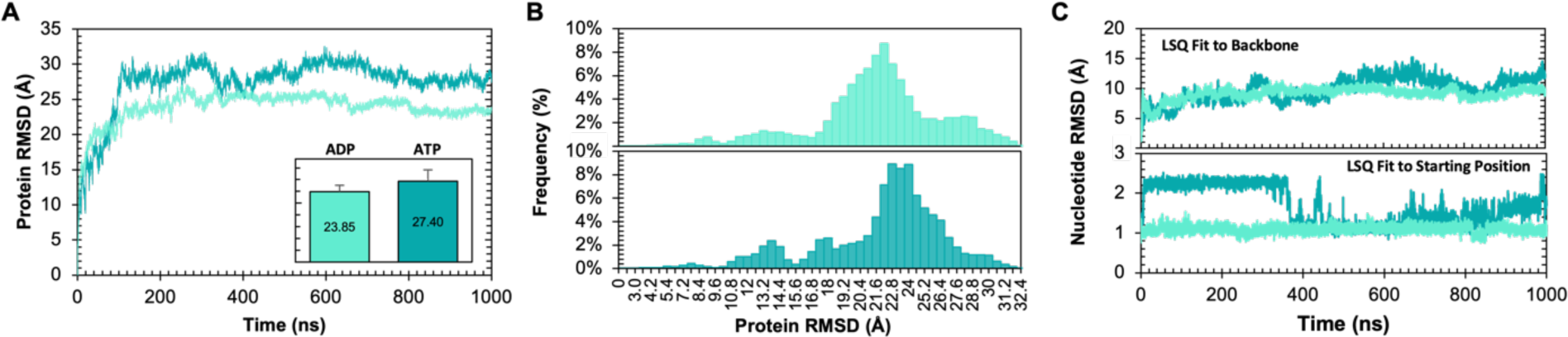
Molecular Dynamics Trajectories Analyses: Protein and Nucleotide RMSD. In (**A**) mean root-mean-square deviation (RMSD) in angstroms (Å) of main-chain protein atoms (N, Cα, C, O), where NLRP31′PYD-ADP is coloured in light teal, and NLRP31′PYD-ATP is coloured in dark teal. RMSD was calculated following a least-squares (LSQ) fit to the equilibrated reference structure at time 0 and plotted as a function of time (0-1000ns). Inset shows the mean ± SD RMSD values for ADP and ATP, with an identical axis range as encompassing plot. In (**B**), histograms of RMSD values for ADP (top) and ATP (bottom) as shown in panel A. In (**C**), nucleotide RMSD values in Å as a function of time in ns, calculated following LSQ to the protein backbone (top) or the starting nucleotide position in the equilibrated structure at time 0 (bottom).

The RMSD values of the nucleotides were calculated relative to the starting position of the protein backbone and the bound nucleotide (**Figure 2C**). The RMSD values, as compared to the equilibrated protein backbone, suggest that both nucleotides were stable within the nucleotide-binding pocket and did not undergo large migrations or artefacts relative to the protein. The position of the ATP nucleotide relative to the starting position of the nucleotide was substantially altered at approximately 400 ns, corresponding to an observed change in the holistic protein structure. This suggests a displacement of the nucleotide-binding pocket due to conformational rearrangements occurring around that time point.

### Stability and Conformations of ADP- and ATP-bound NLRP31′PYD

The solvent-exposed surface area (SASA) of the protein atoms was calculated to evaluate the stability and conformation of the models. The ADP-bound state had fewer solvent-exposed residues over the course of the simulations (**Figure 3A,C**), establishing the expected ‘closed’ conformation for the ADP-bound as compared with ATP-bound protein. This was corroborated by the radius of gyration (Rg) of the protein atoms (**Figure 3B,D**), a measure of the compactness of a system expected to be essentially constant for a protein with a stable tertiary structure. In this case, the ADP-bound system was mainly stable following a small peak and return to baseline during trajectory convergence (prior to ∼30 ns). Conversely, the Rg of the ATP-bound protein increased steadily throughout the simulations. This suggests an opening mechanism in the structure driven by ATP bound within the NACHT domain.

**Figure 3.**
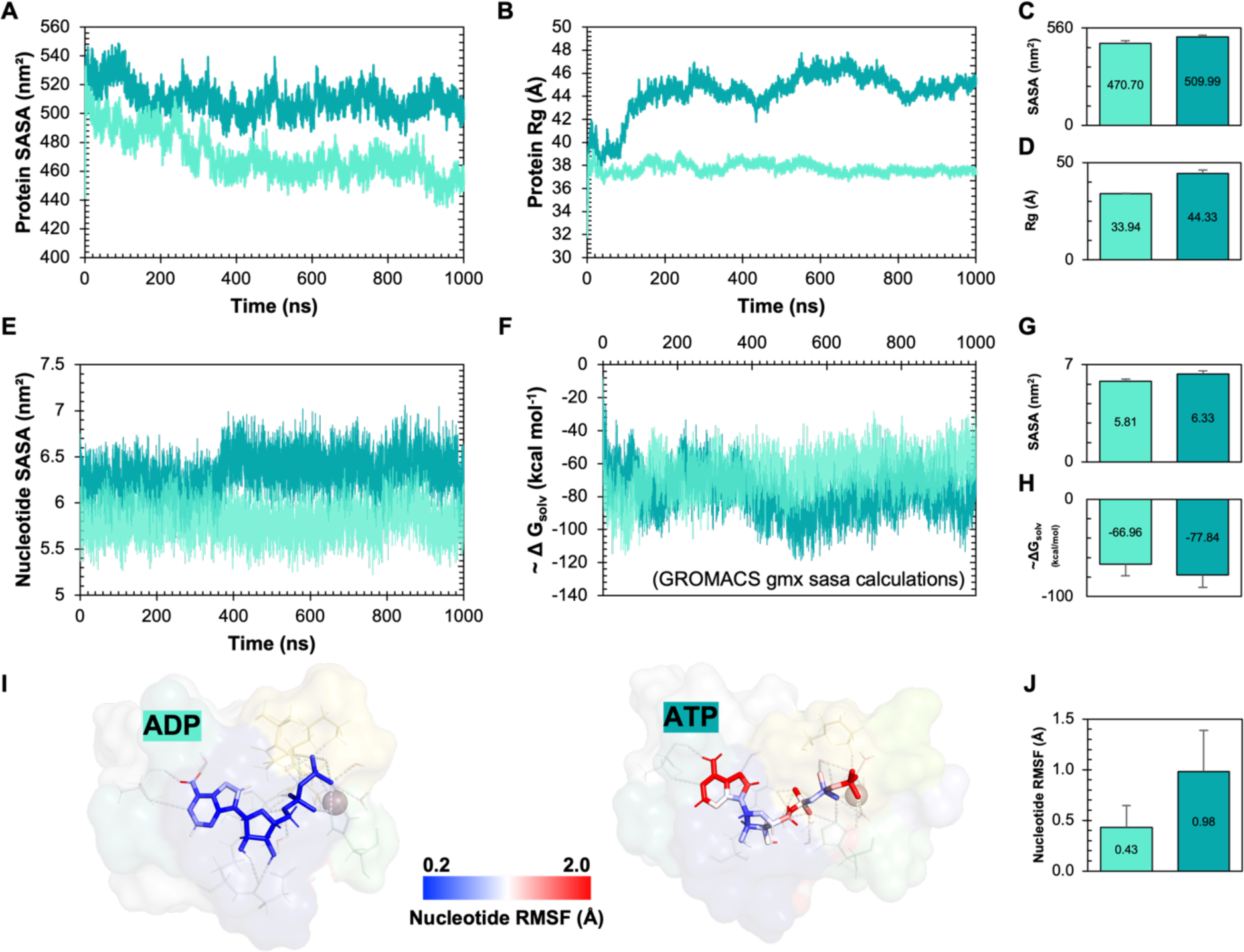
Molecular Dynamics Trajectories Analyses: Solvent Exposed Surface Area, Radius of Gyration and RMSF. In (**A**) and (**B**), the solvent accessible surface area (SASA) and the radius of gyration (Rg, in Å) of all protein atoms in nm^2^ are provided as a function of time (0-1000 ns) for both the ADP- and ATP-bound trajectories. In (**C**) and (**D**), the mean ± SD for SASA and Rg values are provided, respectively. In (**E**) and (**F**), the SASA and GROMACS estimated Gibbs free energy of solvation (ΔG_solv_, kJ mol^−1^) for the ADP and ATP nucleotides are plotted a function of time. Larger values (i.e., less negative) suggest increased stability. In (**G**) and (**H**), mean ± SD for the SASA and ΔG_solv_ are provided for data in panels H and I, respectively. In (**I**), the root-mean square fluctuations (RMSF)of the nucleotide atoms are shown graphically as sticks on a background surface representation of the nucleotide-binding pocket. Nucleotide atoms are coloured in a spectrum from 0.2 Å (blue) to 2 Å (red), as indicated. In (**J**), the mean ± SD RMSF values determined for ADP and ATP are plotted.

Similarly, the nucleotide SASA was increased for ATP relative to ADP (**Figure 3E,G**), where the time course indicated an increase in SASA as the protein structure underwent the opening mechanism. The Gibbs free energy of solvation (ΔG_solv_) was estimated from per-atom solvation energies as a function of exposed surface area. Congruent with the previous data, ΔG_solv_ for ATP-bound NLRP31′PYD was more negative than the ADP-bound form, indicating increased solvation as a function of SASA and the thermodynamic favourability of the opening mechanism in ATP-bound NLRP31′PYD (**Figure 3F,H**). Finally, the per-atom RMSF values were calculated for both nucleotides, suggesting the observed increase in flexibility of the ATP molecule was conveyed by flexibility within the NACHT domain (**Figure 3I,J**).

### Essential Dynamics of MD trajectories

Initial trajectory analyses reveal that substantive conformational changes occurred over the MD trajectories, especially for the ATP-bound model. Principal component analysis (PCA) was used to extract the simulated atoms’ concerted displacements and elucidate the major conformational changes adopted over the trajectories (Lange and Grubmuller 2006). Essential Dynamics (ED) of the protein simulations was employed with the PCA to evaluate the distribution of structural conformations along the trajectories for both nucleotide-bound models of NLRP31′PYD (Amadei, Linssen, and Berendsen 1993; Hayward and de Groot 2008). Following diagonalization of the covariance matrix of Cα atomic fluctuations built from the concatenated trajectories, ED provided a set of eigenvectors and eigenvalues by PCA. The trace values of the covariance matrix of Cα atomic fluctuations were 286 and 592 nm^2^ for ADP- and ATP-bound forms, signifying increased structural flexibility with ATP occupancy. As a result, the configurational space covered by NLRP31′PYD-ATP was significantly larger than that covered by the ADP complex along PC1 and PC2.

Eigenvalues are presented as the proportion of total variance as a function of eigenvector index (i.e, Scree plots). The concerted motions depicted by the first few eigenvalues decrease rapidly in amplitude and approach a totality of constrained and more localised fluctuations. In both cases, the majority of conformational variance was captured within the first few eigenvectors (NLRP31′PYD-ADP), **Figure 4A**; NLRP31′PYD-ATP, **Figure 4B**). PCA projections on the first six eigenvectors are also shown in nm as a function of time for each nucleotide (**Figure 4C,D**). The top 6 eigenvectors comprised 85.1% and 87.2% of the total variance for ADP- and ATP-bound forms of NLRP31′PYD, respectively. The cosine content (CC) (Sang, Liu, and Yang 2020) of each PCA projection was used to evaluate the convergence of conformational sampling during the trajectories (**Figure 4A,B**; insets). Importantly, CC values ranging from 0.007 to 0.243 (NLRP31′PYD-ADP) and from 0.015 to 0.331 (NLRP31′PYD-ATP) demonstrate a high degree of sampling convergence achieved in these simulations. An empirically derived threshold designates CC values <0.5 to be representative of sufficient sampling during MD simulations (Maisuradze, Liwo, and Scheraga 2009).

**Figure 4.**
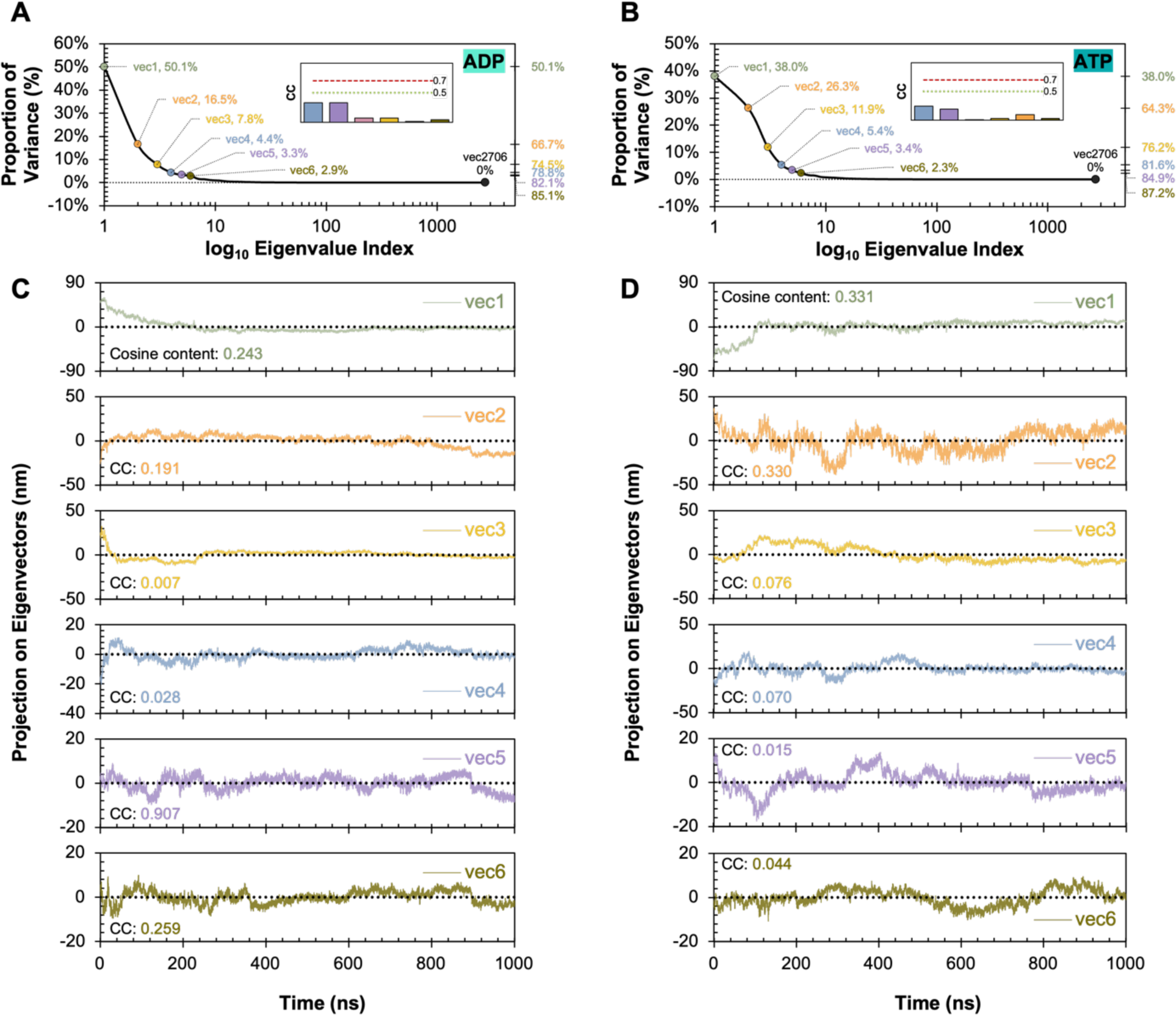
Essential Dynamics Eigenvalue Ranks and Projection Plots with Cosine Coefficients for NLRP31′PYD-ADP and NLRP31′PYD-ATP. Eigenvalue ranks were plotted for NLRP31′PYD-ADP (**A**) and NLRP31′PYD-ATP (**B**). The proportion of variance for each eigen vector is shown as a function of the Eigenvalue Index. Cumulative variance is labeled for the first sex eigenvectors. Inset: the cosine content [CC] was calculated for each eigenvector with the associated validation cut-off values of 0.7 and 0.5 (red and green dashed-lines, respectively). PCA projections (nm) on the first six eigenvectors are shown as a function of simulation time from 0 - 1000 ns for NLRP31′PYD-ADP (**C**) and NLRP31′PYD-ATP (**D**).

Notably, the first eigenvectors (PCs, or classes of motion) contribute 50.1% and 38.0% for the respective NLRP31′PYD-ADP and NLRP31′PYD-ATP models (**Figure 4A** and **4B**), indicating that a significant fraction of the overall atomic fluctuations was captured in these first eigenvectors. Additional inclusion of PC2 and PC3 increased the proportions to 66.7% and 74.5% (NLRP31′PYD-ADP) and to 64.3% and 76.2% (NLRP31′PYD-ATP). Thus, the projections of the first three PCs were evaluated in detail to characterise the collective molecular motions distinguish amongst different conformational sub-states (**Figure 5A-C**). Further evaluation of the per-residue root mean square fluctuations (RMSF) of the first six eigenvectors confirm the major dynamics of each domain and subdomain were captured within the first three PCs for both ADP- and ATP-bound NLRP31′PYD (**Figure 6**). The trace values of the covariance matrix of Cα atomic fluctuations over the 2000 ns simulation were 286 nm^2^ and 592 nm^2^ for NLRP3-ΔPYD(ADP) and NLRP31′PYD-ATP, respectively. So, the results signify that ATP-bound NLRP31′PYD experienced an increased amplitude of atomic fluctuations throughout the simulations with greater overall flexibility and configurational space for PC1 and PC2 as compared to ADP-bound NLRP31′PYD (**Figure 5A,B**).

**Figure 5.**
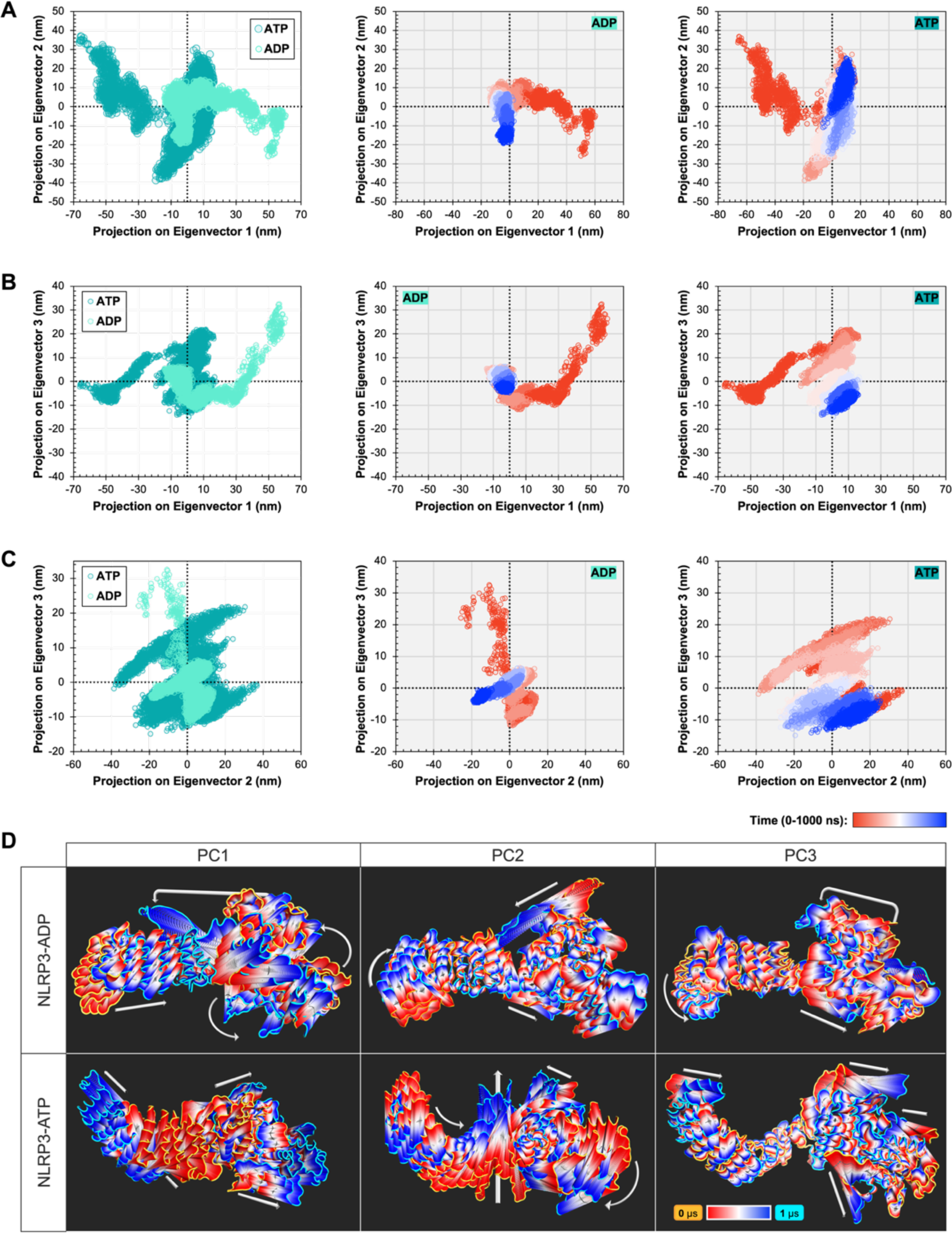
2D-projections of Principal Components with Structural Projections of the Top 3 Eigenvectors. Two dimensional (2D) projections are provided for PC2 vs. PC1 (**A**), PC3 vs. PC1 (**B**), and PC3 vs. PC2 (**C**) throughout the MD simulations. The ADP-(light teal) and ATP-(dark teal) bound NLRP31′PYD conformational states are overlayed in the left-hand column whereas the individual 2D projections for ADP and ATP-bound states are provided in the center and right-hand columns, respectively. The projections are coloured as a function of time (0-1000 ns) as indicated in the heat bar. In (**D**), structural projections along PC1-PC3 eigenvectors for NLRP31′PYD-ADP (top) and NLRP31′PYD-ATP (bottom) are shown for all Cα atoms in NRLP3. Representative structures (50) along the trajectories are coloured according to the time progression. The first and last structures are identified in orange and cyan, respectively. Arrows indicate regions which undergo the largest conformational changes within each projection.

**Figure 6.**
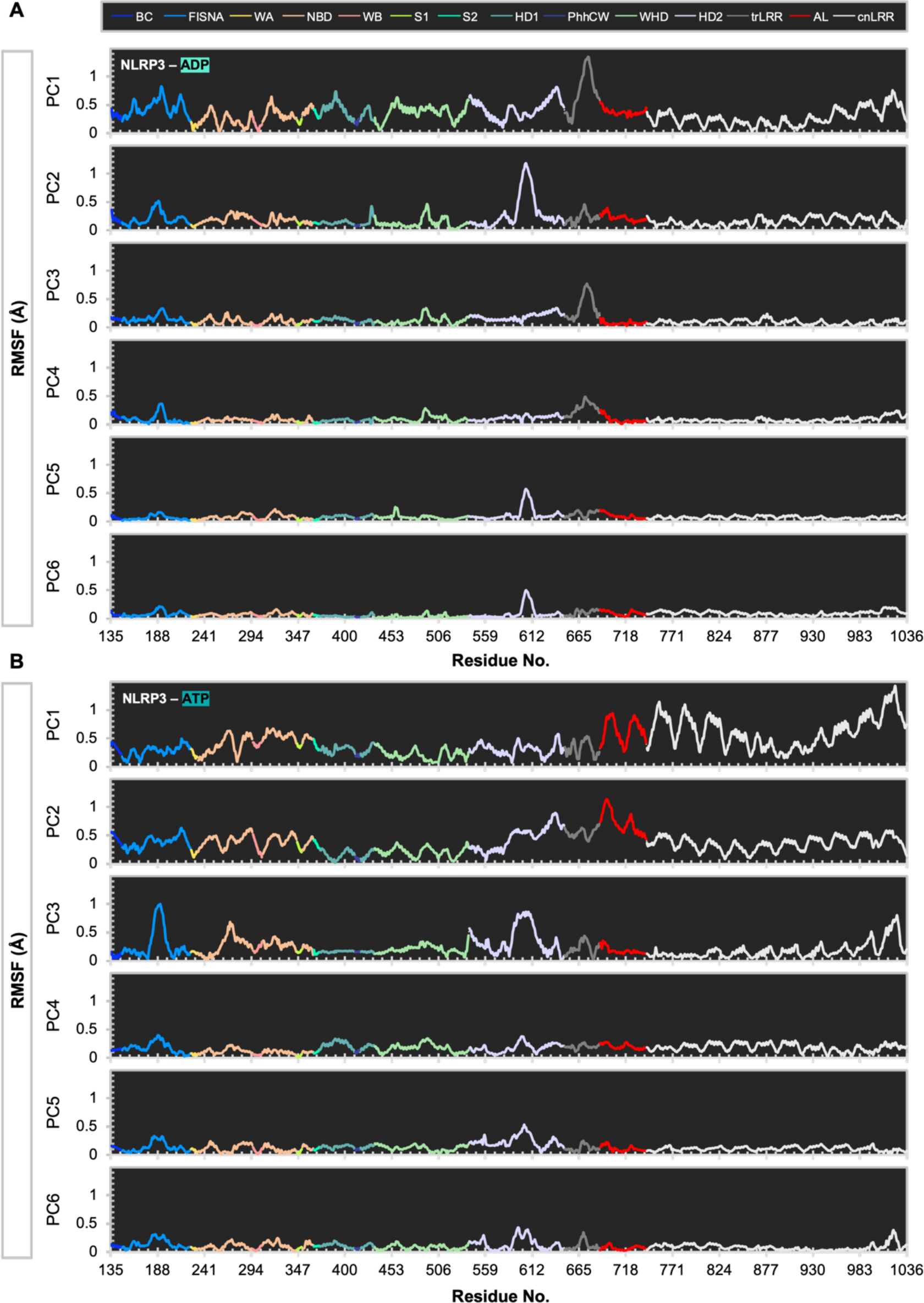
Cα Root Mean Square Fluctuations (RMSF) per Residue of Eigenvectors 1-6 for NLRP31′PYD. Data are provided for NLRP31′PYD-ADP (**A**) and NLRP31′PYD-ATP (**B**); colouring is by domain and motif as indicated.

### Free energy landscape (FEL) of NLRP3 Configurational Spaces

The temporal path of the trajectories connecting free-energy minima and the sub-conformational spaces adopted by the NLRP31′PYD complexes are displayed in **Figure 7A**. In general, the trajectories connecting the minima suggest a more progressive and unambiguous path for NLRP31′PYD-ADP as compared with NLRP31′PYD-ATP. Numerous distinct configurations of NLRP31′PYD-ATP within each energy bin are suggested. The free energy landscape (FEL) plots suggest distinct configurations in each grouping. ADP-bound NLRP31′PYD adopted several conformations during equilibration (**Figure 7B**) and reached an energy minimum first at 173.2 ns, fluctuating to a higher energy state (>1.86-3.71 kcal mol^−1^) before returning to the minimum energy at 239.9 ns (**Figure 7C**). Following energy fluctuations between 243 and approximately 285 ns, a low energy conformation was adopted and remained relatively consistent until 660 ns, when energy values fluctuated more rapidly until the secondary lower energy states were achieved. Additional basins indicate supplementary low-energy structures adopted as the trajectories progressed to 1000 ns, though the first structure presents the deepest energy basin (i.e., lowest ΔG_solv_). From these data, it was concluded that the trajectory converged in less than 300 ns, and a second low-energy conformational state of NLRP31′PYD-ADP was visited towards the end of the trajectory. ATP-bound NLRP31′PYD adopted more conformational states than NLRP31′PYD-ADP during the equilibration (**Figure 7D**), yet quickly converged on the energy minimum within 107 ns (**Figure 7E**). Overall, the ED and FEL data indicate notable distinctions in atomic fluctuations, conformational status, and dynamic behaviour as a direct result of bound nucleotide identity for NLRP3. The 3D energy plots, in particular, highlight the numerous conformations adopted by ATP-bound NLRP31′PYD, while the energy by time plots indicates relatively low (i.e., favourable) free energies associated with that flexibility. These data also support the ΔG_solv_ data in which the ATP structure was more thermodynamically favourable, more flexible, and able to adopt more conformational states than ADP-bound NLRP31′PYD.

**Figure 7.**
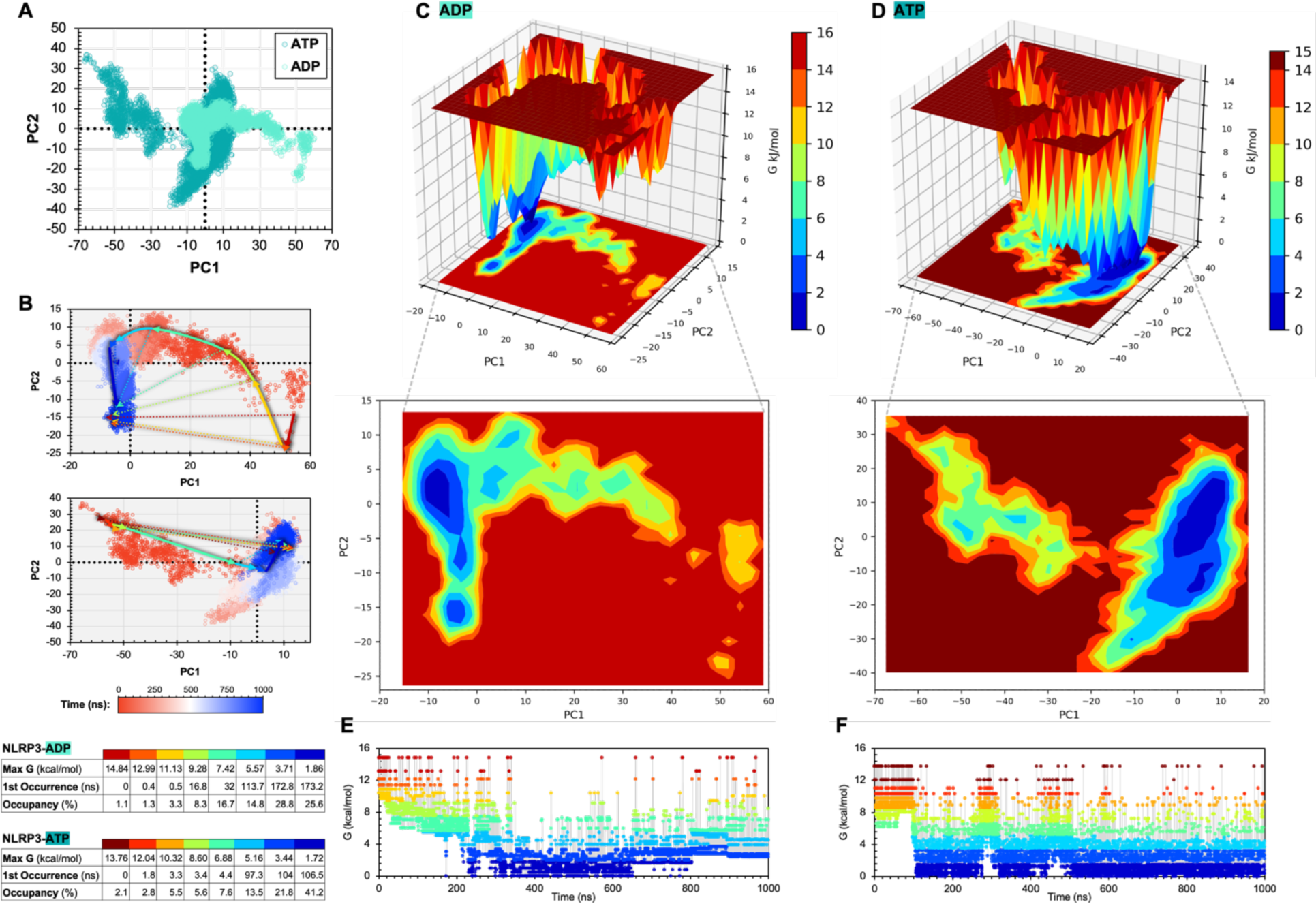
Essential Dynamics of NLRP31′PYD-ADP and NLRP31′PYD-ATP Trajectories. In (**A**), ADP (*i*) and ATP (*ii*) plots are separated and coloured as a function of time (0-1000 ns) as indicated. Solid arrows were drawn from the minima of each energy basin to demonstrate the energy course over the trajectory. Dotted arrows indicate the first frame to the final frame within the basin, highlighting the overall spread of each bin to final minima within each free energy grouping. Arrows are coloured to align with Gibbs free energy (ΔG, kJ mol^−1^) as in panels (**B**) and (**C**). The tables (*iii*) indicate the first occurrence of the energy basin (ns) and the occupancy (%) of the energy status over the trajectory. The free energy landscape (FEL) analyses of ADP- and ATP-bound NLRP31′PYD in 3D (**B** and **C**) and 2D (**D** and **E**) for PC1 and PC2. FEL plots were generated with modified python scripts [fel3d.py and fel2d.py]. Colour bars represent the ΔG, (kJ mol^−1^) energies of the conformational states adopted throughout the trajectories, where low-energy minima are in blue, and the highest energy conformational states are in red. In (**F**) and (**G**), ΔG is plotted as a function of simulation time for ADP- and ATP-bound NLRP31′PYD, respectively. PCA and free energies were assessed with the in-built GROMACS functions gmx covar, gmx anaeig and gmx sham.

### Holistic fluctuations in NLRP31′PYD structure with ADP/ATP-binding

The RMSF about the average position of residues defined for backbone Cα-atoms was used to quantify the per-residue dynamics in ADP- and ATP-bound NLRP31′PYD (**Figure 8A**). Fluctuations in NLRP3-ATP were higher at nearly every residue, supporting the RMSD data that ATP occupancy resulted in a more flexible structure. The largest overall deviations between nucleotide-binding models were observed in the LRR domain with a ΔATP/ADP of 4.72 Å (**Figure 8B**). In descending order, the remaining ΔATP/ADP RMSF values in Å were 4.45 (S2), 4.18 (HD2), 3.81 (FISNA), 2.91 (S1), 2.67 (WHD), 2.54 (WB), 2.51 (HD1), 2.38 (WA) and 1.08 (PhhCW). The S2, S1, WB, WA and PhhCW motifs were all directly involved in stabilising the bound nucleotides (**Figure 8C**), and thus the deviations in these regions are logical due to differential nucleotide interactions with those motifs.

**Figure 8.**
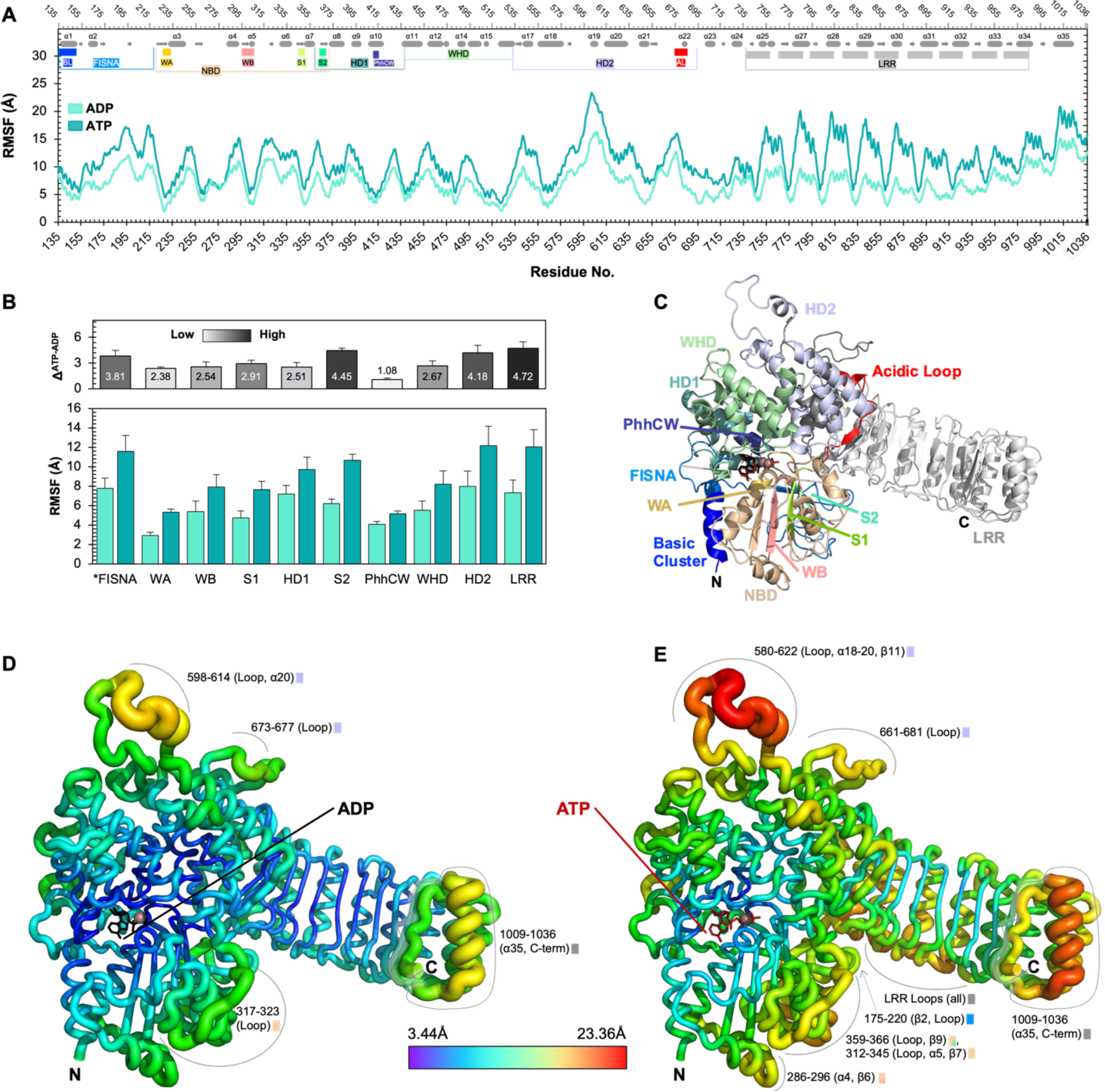
Root Mean Square Fluctuations (RMSF) in ADP- and ATP-Bound NLRP31′PYD Structures. In (**A**), RMSF (Å) as a function of NLRP3 residues from 135 to 1036. Secondary structure elements are shown at the top as rounded rectangles (α-helices) or arrows (β-sheets), with domains, subdomains and motifs coloured as in panel (C). In (**B**), RMSF values (mean ± SD) of various domains and motifs are shown for both the ADP- and ATP-bound NLRP31′PYD trajectories. (Top), ΔRMSF (ATP-ADP) values are shown for each domain/motif and coloured in a spectrum as indicated (3.4 to 23.4 Å), with the largest value in black and the lowest in white. In (**C**), cartoon representation of nucleotide interaction domains/motifs for NLRP31′PYD, coloured as labelled. In (**D**) and (**E**), comparisons of the structural flexibility of ADP- and ATP-bound NLRP31′PYD protein backbones, respectively. Per-residue average backbone RMSF values are mapped on the starting structures, where thicker tubes denote higher RMSF and are coloured in a rainbow spectrum as indicated. Key regions with high relative RMSF are labelled with associated secondary structure and a legend marker corresponding to the domains/motifs coloured in panel A (top) and the inset figure of panels C and D. All RMSF values were calculated from two separate MD simulations (1000 ns) for each structure.

Interestingly, the ΔATP/ADP fluctuations in all motifs apart from S2 (located within the HD1 domain) were smaller than those of the LRR, HD2, FISNA and WHD domains. These observations suggest that direct interactions with the nucleotide convey a global structural reorganisation that was not restricted to local conformational changes. Of particular interest is the HD2 domain, located between the NACHT and LRR domains, where a hinge mechanism was noted in the ATP-bound structure over the course of the trajectories. In particular, the unstructured loop between α-helices 18 and 20 and α-helix 35 in the final LRR experienced maximal residual fluctuations in the ATP-bound state, contrary to the ADP-bound state (**Figure 8D,E**). This highlights the importance of the HD2 hinge region in ATP-mediated activation, previously suggested to mediate intracellular K^+^ sensing and conformational changes leading to activation (Tapia-Abellan et al. 2021). In the structure of full-length NLRP3 (PDB: 7PZC), the FISNA domain interacts heavily with the core NBD to stabilise the ADP-bound structure (Hochheiser et al. 2022). Even with four FISNA residues missing from these models, RMSF was increased for NLRP31′PYD with ATP occupancy.

### GROMOS Clustering of the Conformational Trajectories

Concatenated replica trajectories were clustered by geometric similarity using the GROMOS algorithm (Daura et al. 1999) to explore the variety of protein conformations adopted in each nucleotide-bound state. In this regard, GROMOS analysis resulted in an increased number of individual clusters, quantitatively indicating that ADP conformations adopted over the trajectories were less dissimilar than those resulting from ATP binding (**Figure 9A**). Cluster IDs (a total of 6 and 9 for ADP and ATP, respectively) were identified at a cut-off of 0.9 nm within the simulation time course provided in **Figure 9B,C**. The largest clusters in the ADP- and ATP-bound NLRP31′PYD runs (i.e., clusters 5 and 7, respectively) demonstrate the structural discrepancies elicited by the bound nucleotide. The ADP-bound structure was more compact and stable, leading to differential hinging between the LRR and NACHT domain in the two models. Specifically, the ATP-bound NACHT domain underwent an opening mechanism associated with hinging at the NACHT-HD2-LRR axis. This protein motion resulted in an outward projection of the HD2 and LRR domains and an opening of the NACHT domain (**Video 1**).

**Figure 9.**
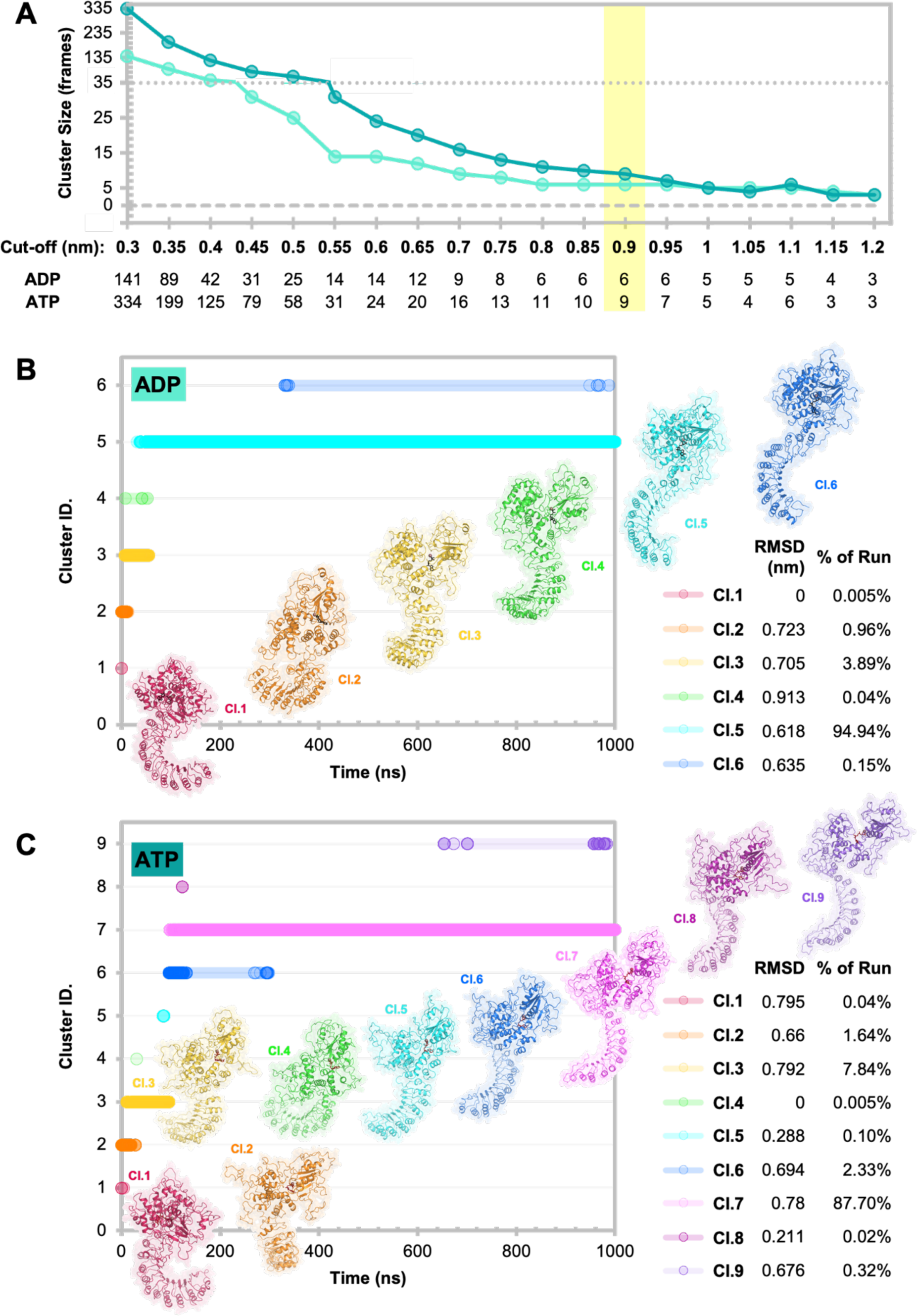
GROMOS Clustering of ADP- and ATP-bound NLRP31′PYD Trajectories. Two 1000 ns trajectories for the ADP- and ATP-bound NLRP31′PYD structures were concatenated and analysed for clusters with RMSD cut-off values highlighted. In (**A**), cluster size (i.e., number of frames within the RMSD cut-off) for each MD trajectory was plotted against the RMSD cut-off (nm) with associated data provided for ADP- and ATP-bound forms. The 0.9 nm RMSD cut-off was selected and highlighted in yellow, where six (NLRP31′PYD-ADP) and nine (NLRP31′PYD -ATP) medoid structures represent the conformations observed during the trajectories. Note the split vertical axis represented by the dotted line with 0 shown as a thicker dashed line. In (**B**) and (**C**) cluster IDs are plotted as a function of time, for NLRP31′PYD-ADP and NLRP31′PYD-ATP, respectively; the IDs represent individual frames adopting the conformation of each cluster. Clusters are sorted by the first occurrence. The RMSD of all structures within each cluster (Cl) is shown in a table with the legend, alongside the percentage of each cluster observed in the trajectory. RMSD values of 0 represent clusters containing only 1 frame. Medoid structures for each cluster are displayed as cartoons with transparent surfaces, with sticks representing ADP (in black) and ATP (in dark red). Mg^2+^ atoms are shown as violet spheres. Medoid representations are individually labelled by cluster.

### Geometric Volumes and Displacement of Medoid NACHT Domains

To further characterise the ‘opening’ mechanism observed for NLRP3 during ATP binding, associated volumes and areas for the NACHT domain of the medoid structures of ADP (cluster 5) and ATP (cluster 7) were calculated (Chen and Makhatadze 2017). As shown in **Figure 10A**, the ADP-bound NACHT domain had a molecular surface area of 38,556 Å^2^ with a solvent-exposed surface area of 13,686 Å^2^. The surface area of the NACHT domain of the largest ATP-bound cluster was increased only slightly (∼3%) to 39,744 Å^2^, but with a much larger (∼53%) increase in the SASA to 21,004.26 Å^2^. Additionally, the total volume and van der Waals (vdW) volumes as well as packing density (i.e., the ratio of vdW volume to solvent excluded total volume) establish the increased atomic packing of the NACHT domain with ADP occupancy.

**Figure 10.**
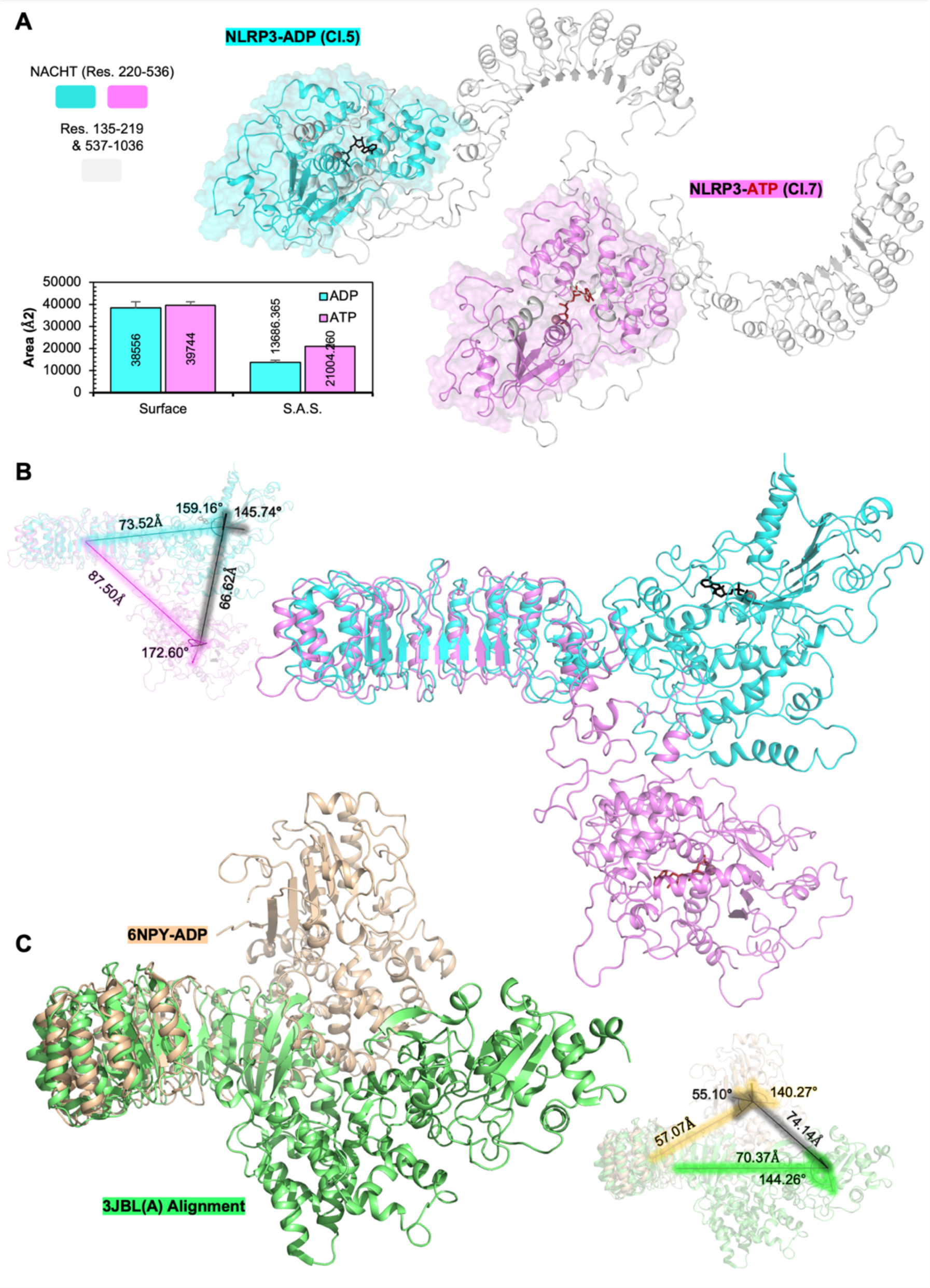
Conformational Displacement of ADP-vs. ATP-bound NLRP31′PYD Medoid Structures. In (**A**), the NACHT domains of the largest cluster medoid structures of ADP-(cyan) and ATP-(pink) bound NLRP31′PYD are shown in a surface representation, with other residues in white. Total areas and SASA are listed and plotted in the inset bar chart (n=2 ± SD). In (**B**), the NLRP31′PYD-ADP and -ATP structures are aligned by the LRR domain. The inset figure highlights the distances of displacement in angstroms (Å) and the angle (°) between the NACHT and LRR domains. The black displacement vector and angle were calculated between NACHT domains. In (**C**), The first chains [A] of inactive NLRP3 (PDB ID: 6NPY) and active NLRC4 (PDB ID: 3JBL) are aligned as in panel B. NACHT and LRR domains of MD simulation medoid structures used for calculations included residues 220-536 and 699-1036, respectively. The domain boundaries for the inactive 6NPY structure and the active NLRC4 structures were 218-534 and 93(N-terminus)-222 for the NACHT, and 697-1036(C-terminus) and 489-1024(C-terminus) for the LRR, respectively. Angles and displacement vectors were calculated in Pymol with a script obtained from the open-source Pymol repository [orientations.py]. Areas were computed with Protein Volume 1.3.

The angles between the NACHT and LRR domains of the largest NLRP31′PYD-ADP and NLRP31′PYD-ATP cluster medoid structures were quite distinct, with angles of rotation of 159.2° and 172.6° for the ADP and ATP bound structures, respectively, and concomitant displacements of 73.5 Å and 87.5 Å (**Figure 10B**, 2.0% and 19.0% increases for the ATP model). Interestingly, these data are consistent with the ratios observed when comparing the ADP-bound cryo-EM structure of NLRP3 (PDB: 6NPY; chain A) with the NLRP31′PYD-ATP structure aligned to the activated NLRC4 structure (6NPY alignment to 3JBL; chain A), where the activated/inactive NACHT-LRR angle ratio was 2.8%, and the displacement ratio was 23.3% (**Figure 10C**).

### Conformational Changes are Regulated by Variable Nucleotide Interactions

To identify the interactions within the nucleotide-binding site which drive the global distinctions between models, Molecular Mechanics Poisson–Boltzmann and Surface Area continuum solvation (MM-PBSA) binding energies, protein-nucleotide hydrogen bonds and internal hydrogen bonds were calculated for each trajectory. MM-PBSA calculations reveal distinctions in binding affinity between ADP and ATP, and variable per-residue ligand interaction energies elucidated critical residues in the differential modes of ATP and ADP binding by NLRP3. Non-bonded electrostatic terms (Δ_ele_) contributed the most to considerable discrepancies between models, underscoring the energetic favourability of the additional negative charge carried by the ATP molecule in binding to NLRP31′PYD (**Table 1**). Additionally, extensive polar contributions to the binding of both nucleotides were noteworthy, with increased solvent accessible areas resulting in positive ΔE_PB_ values for both models, increasing from 316.1 ± 3.6 for ADP binding to 562.5 ± 10.2 kcal mol^−1^ for ATP binding. These were compensated by highly negative entropies (TΔS) calculated from the trajectories, resulting in negative (favourable) binding interactions for both nucleotides. A considerably lower binding affinity of NLRP31′PYD for ADP was calculated, with estimated ΔG_bind_ of -8.7 ± 3.9 kcal mol^−1^ while ATP demonstrated a high affinity for NLRP3 (ΔG_bind_ = -62.5 ± 24.6). Interaction energy standard deviations (σIE) for both complexes fell beneath an established validation cut-off of 3.6 kcal mol^−1^ (Ekberg and Ryde 2021), with values of 0.9 and 3.10 for ADP and ATP, respectively. Additionally, entropy results fell within the expected values for estimation by IE (Ekberg and Ryde 2021).

**Table 1.**
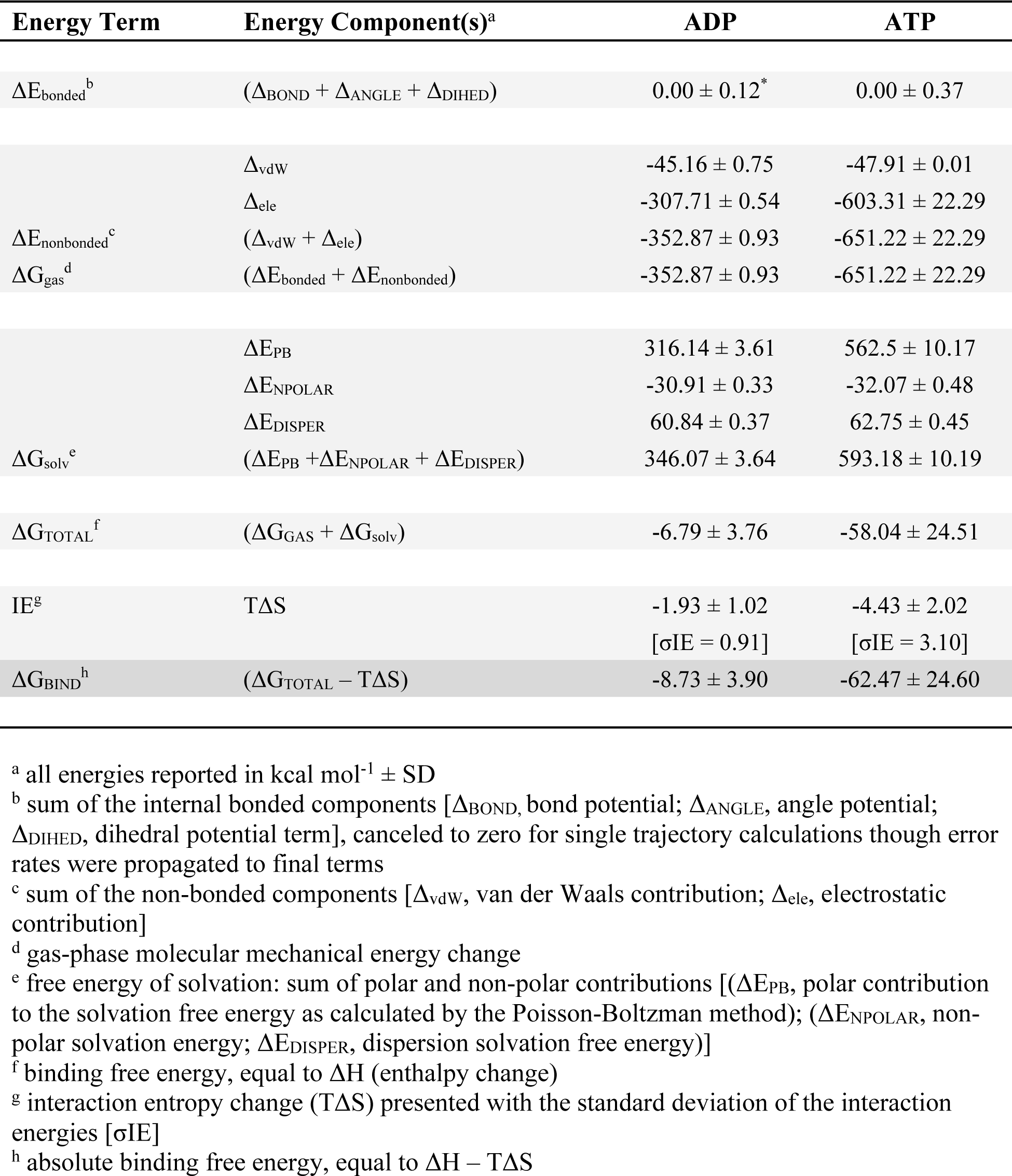
MM-PBSA ligand binding free energies of NLRP31′PYD.

Overall, small error rates and low σIE values indicate accurate energy estimates whereby ATP binds NLRP31′PYD with an over 7-fold increase in affinity compared with ADP. These data indicate not only increased binding affinity for ATP as compared with ADP, but also shed light on the mechanisms involved in ATP binding. In this case, ATP binding in the active site resulted in higher solvation energies (ΔG_solv_) which were compensated by a highly negative entropic contribution as the protein underwent global rearrangement. In short, ATP binds the NACHT with high affinity and drives conformational rearrangement to higher energy states, which are by and large more “open” or solvent accessible than those adopted by NLRP31′PYD-ADP. Finally, NLRP31′PYD-ADP demonstrated increased internal hydrogen bonding throughout the simulation, 600 ± 20 compared with 575 ± 16 for NLRP3-ΔPYD(ATP). This supports the previous findings and suggests increased stability for ADP-bound NLRP31′PYD compared to NLRP31′PYD-ATP.

### Per-residue Energy Decomposition Analysis of the NLRP31′PYD-Nucleotide Complex

The binding energies from residues within 6 Å of the nucleotide were calculated with the MM-PBSA method and are shown along with hydrogen bond occupancies in **Table 2**. Residues with the largest changes in absolute decomposition values (ΔATP-ADP) are presented in **Figure 11**. Several residues were predicted to contribute substantively to ADP binding: including Asp153, Arg154, and Asn155, Tyr168 and Thr169 within FISNA, Ile230 and Gly231 within WA, along with Pro412 and Leu413 within PhhCW. Residues with greater contributions to ATP binding include Gly229, Lys232, Thr233, and Ile234 within WA, Asp302 and Gly303 within WB, Tyr381 in HD1, as well as His522 in WHD. In both cases, the dominant contribution to nucleotide-binding was associated with residues within the WA. Several residues (i.e., Ala228, Gly229, Ile230, Gly231, and Lys232) contributed favourable individual binding energies ranging from -1.7 to -5.9 kcal mol^−1^ for ADP and from -4.5 to -6.1 kcal mol^−1^ for ATP. An exception was Ala227, which exhibited the least energetically favourable value for NLRP31′PYD binding of ATP. Only Pro412 was less favourable than Ala227 in the process of ADP binding. The Gly226 residue, while located within the WA, did not fall under the distance cut-off for ADP so was ranked as one of the least favourable residues for ATP binding.

**Figure 11.**
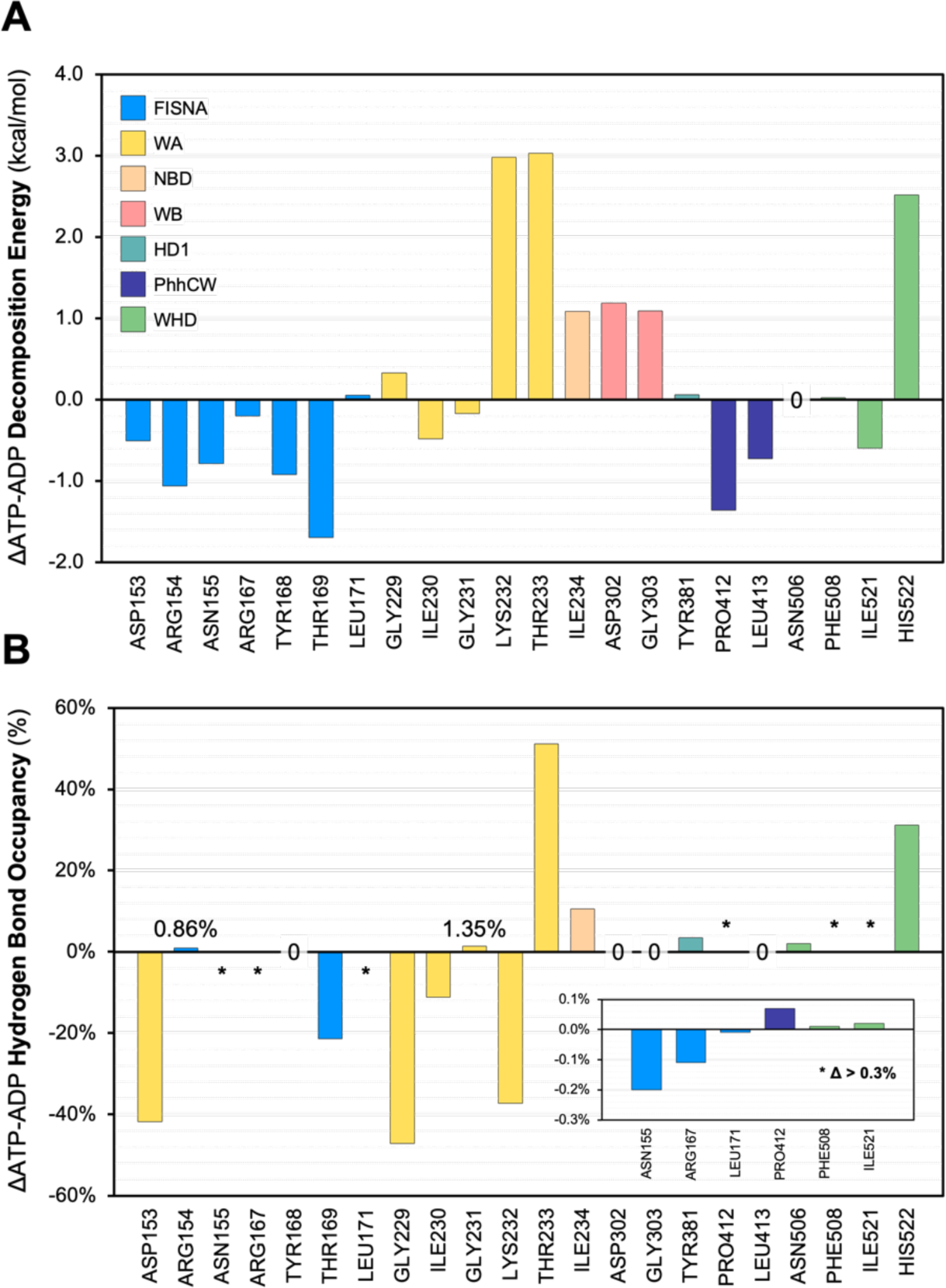
1′ATP-ADP Decomposition Energies and Hydrogen Bond Occupancies. Residues with either absolute ΔATP-ADP decomposition energies > 0.7 kcal/mol and any hydrogen bonding over the trajectories are shown in panels (**A**) and (**B**), respectively. Absolute values were used to calculate the ΔATP-ADP for decomposition energies. Thus, in both panels, positive values indicate a larger contribution to NLRP31′PYD-ATP binding, while negative values indicate a larger contribution to NLRP31′PYD-ADP binding. Note that positive or negative values do not indicate energetic favourability or otherwise, see Table X for true values. Colour coding for the domains is presented in the first panel. Δ values equal to zero are labelled as such. In the inset panel provided for (B), residues with Δ hydrogen bond occupancy values >0.3% (labelled with an asterisk) are shown on a small axis range from -0.3 to 0.1%.

**Table 2.**
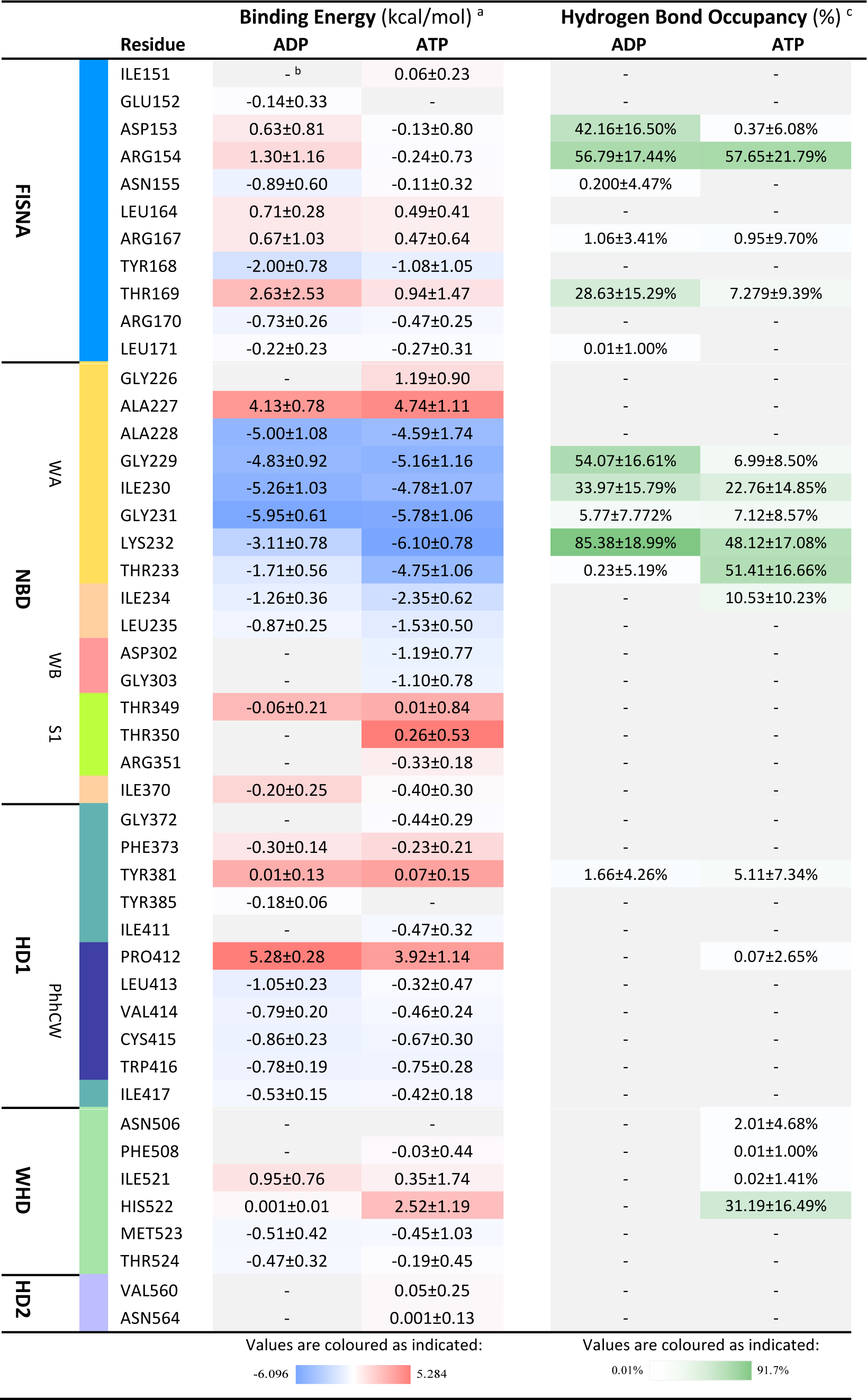
Per-residue MM-PBSA binding energies and hydrogen bond occupancies.

Highly unfavourable interaction energies were identified with ADP occupancy for Thr169 (FISNA), Pro412 (PhhCW) and His522 (WHD) but thermodynamic improvement was observed with a shift to lower positive values upon ATP binding (**Table 2**, **Figure 11**). Charged FISNA residues Asp153 and Arg154 were unfavourable contributors to ADP binding, with a shift to mildly favourable contributions during ATP binding. Additionally, favourable energy contributions from Lys232 and Asp302 were increased substantially with replacement of ADP for ATP binding. Taken together, the results suggest that the binding energies of all charged residues within the active site were more favourable with ATP when compared to ADP. These residues account for the large distinction in electrostatic favourability for ADP and ATP binding. Notably, His522 was a negligible contributor to ADP binding yet a relatively large unfavourable contributor to ATP binding. So, the opening mechanism observed within the NBD that was translated to the global structure was presumably influenced by this repulsive effect. Finally, favourable decomposition energies (i.e., less than -1 kcal mol^−1^ for both ADP and ATP) were observed for Tyr168 (FISNA), the previously discussed WA residues, and Ile234 (NBD). Notably, Asp302 and Gly303 of the WB were significant contributors to ATP binding only, falling above the distance cut-off (6 Å) for calculation in the ADP-bound model.

### ADP/ATP Hydrogen Bonding in NLRP31′PYD

Hydrogen bond occupancies (**Figure 12**) were calculated for all residues engaged with ADP or ATP (i.e., when the donor and acceptor atoms from the NLRP31′PYD protein and the nucleotide were within 3 Å with an angle cut-off of 20° over 20,000 calculated frames). Hydrogen bonding events were also determined from the complete 1000 ns trajectories and are listed in **Table 2**. The number of protein-nucleotide hydrogen bonds was greater for NLRP31′PYD-ADP than for NLRP31′PYD-ATP, with the mean number of bonds over the trajectories calculated to be 3.1 ± 1.2 and 1.9 ± 1.1, respectively. Per frame hydrogen bonds ranged from 0 to 7 (ADP) and 0 to 6 (ATP), and the number of unique residues interacting with the nucleotide ranged from 0 to 7 in both cases. The distribution of bonds per frame and the unique number of bonding residues per frame are provided in **Table 3**. While the total number of residues within the NBD shown to provide hydrogen bonds with ATP increased to 16 from 12 in ADP binding (**Figure 12**), the number of bonds and unique residues per frame decreased as compared with ADP, indicating a more variable topology for the nucleotide-binding pocket over time.

**Figure 12.**
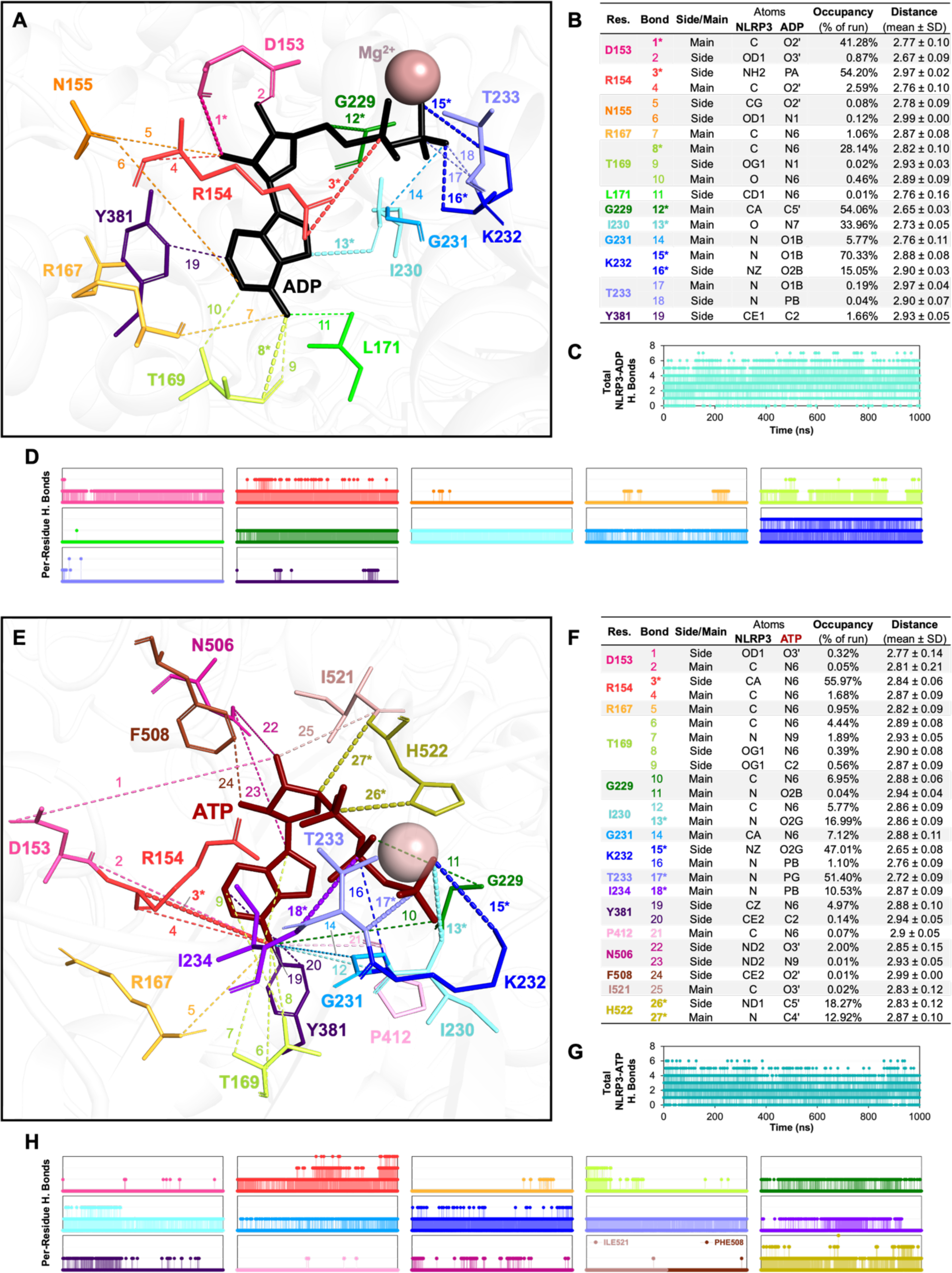
Hydrogen Bonding of NLRP31′PYD with Nucleotides. NLRP31′PYD hydrogen bonds to ADP (**A-D**) and ATP (**E-H**) were calculated over all frames for two separate MD simulations with the HBonds_VMD plugin. Hydrogen bonds were defined between two polar atoms less than 3 Å apart with an angle cut-off of 20°. Bonds are presented graphically in (**A**) and (**E**), where bonds and residues are coloured in a rainbow spectrum, and colours are held constant between both panels. Exceptions include residues binding to only ADP (N155, Y168, L171 and W416) or only ATP (I234, P412, N506, F508, I521 and H522). Data in (**B**) and (**F**) include bond numbers, colours and other details, including whether the bonding occurs from the main or side chain of the residue, names of the protein and nucleotide atoms, mean occupancy of the bond as a percentage of the total simulation time and mean distance (Å) between the donor and acceptor atoms involved in each bond. Hydrogen bonds present for over 10% of the total concatenated trajectories are shown graphically as thicker dashes, and bond numbers are labelled with an asterisk. In (**C**) and (**G**), the mean total number of hydrogen bonds as a function of simulation time (1 μs) are represented for ADP and ATP, respectively. Bonds are deconstructed by individual NLRP3 residue in panels (**D**) and (**H**), where all bonds to each residue are represented in the same colour as shown in the graphics and data tables. Axes run from 0-1000 ns on the abscissa and 0-3 hydrogen bonds on the ordinate. Note that in (H), the bonds from Ile521 and Phe508 to ATP are shown in a single panel due to space constraints and non-overlapping bonds during the trajectories.

**Table 3.** Per-frame number of Hydrogen Bonds and Unique Bonding Residues.

| Bonds /<br>Frame <sup>a</sup> | ADP |  | ATP |  | Residues<br>/ Frame <sup>b</sup> | ADP |  | ATP |  |
| --- | --- | --- | --- | --- | --- | --- | --- | --- | --- |
|  | H Bonds | % of Total | H Bonds | % of Total |  | Bonded Res. | % of Total | Bonded Res. | % of Total |
| 0-2 | 3160 | 31.6% | 7122 | 71.2% | 0-2 | 6818 | 34.1% | 11002 | 55.0% |
| 2-4 | 5552 | 55.5% | 2709 | 27.1% | 2-4 | 11266 | 56.3% | 8218 | 41.1% |
| 4-6 | 1267 | 12.7% | 171 | 1.7% | 4-6 | 1910 | 9.5% | 782 | 3.9% |
| 6-7 | 23 | 0.23% | 0 | 0% | 6-7 | 10 | 0.05% | 2 | 0% |

<sup>a</sup> Number of hydrogen bonds totaled for each frame in 2×1μs MD simulations; binned in intervals of 2 with an overflow bin for values = 7. Histograms for ADP (teal) and ATP (dark teal) shown below.
<sup>b</sup> Number of unique residues in hydrogen bonding interactions with the nucleotide per frame, binned as in <sup>a</sup>. Histograms shown below.

The stability of the NBD during ADP occupancy is highlighted by the increase in total bonds and stability of specific residue bonds observed over the full trajectories. The increase in hydrogen bonding with ADP occupancy helps to explain the significant decrease in unfavourable polar contributions to binding affinity observed for ADP (i.e., positive ΔE_PB_ values of 316 ± 4 for ADP and 563 ± 10 for ATP, **Table 1**). The increased number of bonding residues in conjunction with decreases in the number of per-frame bonds and residues suggests the opposite for ATP, where stable bond formations were more sporadic. This is illustrated in **Figure 12D** and **12H**, where bonds associated with each residue highlight the transitory nature (<10% occupancy) for 10 of the 16 bonded residues over time (mean values are provided in **Table 2**). These include bonds between the nucleotide and NLRP3 residues Asp153, Arg167, Thr169, Gly229, Gly241, Tyr381, Pro413, Asn506, Phe508 and Ile521. In contrast, some of these residues represent the most stable ADP bonding events, including the FISNA residues Asp153 (occupancy=42%) and Thr169 (29%), along with the WA residue Gly229 (54%) (**Figure 11B**). Additionally, Lys232(WA)-ATP hydrogen bond occupancy increased from 48% to 85% for Lys232-ADP.

Protein-nucleotide hydrogen bonds were not correlated with binding energies; however, residues that display differential energies for ATP or ADP binding were primarily characterised by increased hydrogen bond occupancies to either nucleotide, with three exceptions (**Figure 11B**). Gly229 (WA) demonstrated similarly favourable energetic contributions to ADP and ATP binding, while ATP-hydrogen bonds were formed for only ∼7% of the trajectories and ADP bonds were formed for nearly 55% of the simulations (**Table 2**). The ΔATP-ADP decomposition energy for Lys232 (WA) was more substantial, approximately doubling the favourable binding contribution for ATP compared to ADP. However, the likelihood of hydrogen bonding for Lys232 decreased substantially for ATP, with a ∼48% occupancy compared with ∼85% for ADP. Moreover, the binding energies for Pro412 in the PhhCW motif were unfavourable in both cases, increasing by 1.36 kcal/mol for ADP binding. Concordantly, no hydrogen bonds were detected from Pro412 to ADP, and very few bonds formed with ATP (< 1% occupancy over the trajectory). This suggests a negative repulsion event to both nucleotides, which is comparatively decreased upon ATP binding. This observation offers another explanation for the highly positive ΔE_PB_ value associated with bound ATP. Small increases in hydrogen bond occupancy upon nucleotide change from ADP to ATP were also observed for Arg154 (FISNA), Gly231 (WA), and Tyr381 (HD1), and unique bonds elicited with ATP (i.e., ADP hydrogen bond occupancy = 0) included residues Asn506, Phe508, and Ile521 of the WHD. Moreover, two additional residues were observed to have increased hydrogen bonding to ATP: Thr233 (WA) and His522 (WHD). The large Δ binding energies represent increased favourability for ATP binding compared to ADP (**Figure 11**, **Table 2**). These residues, alongside Lys232, were the predominant favourable contributors to the large difference in calculated binding affinities listed in Overall, these data are indicative of a less flexible NBD in NLRP3ΔPYD-ADP contrasted with NLRP3ΔPYD-ATP, as shown by the increased number of stably bound residues and the total number of hydrogen bonds over time. While increased hydrogen bonding was noteworthy in decreasing the unfavourable polar energetic contributions calculated by MM-PBSA, the highly favourable electrostatic contributions to ATP binding result in a substantially increased affinity for ATP, together with a more flexible NBD). Together with previous data, this reveals the mechanism by which ATP binding results in proximate conformational changes that are conveyed to the global structure through highly favourable electrostatic interactions and decreased unfavourable polar interactions relative to ADP.

### Inter-domain bonding observed throughout the trajectories

The number of internal hydrogen bonds amongst various domains were parsed to investigate differential domain interactions triggered by the bound nucleotides (**Figure 13A**). While no bonds were formed between the NBD and LRR or between the WHD and LRR, the mean number of inter-domain bonds was increased throughout the NLRP3ΔPYD-ADP trajectories in nearly all cases (**Figure 13B**). The overall correlation of the trajectories observed for inter-domain hydrogen bonds highlights the poor conservation of hydrogen bonding patterns which in turn reflect distinctions in global conformation driven by the bound nucleotide. The highest correlated bonding patterns occurred between NBD and WHD as well as between HD1 and HD2. Both increased over the simulation time, although the r values indicated modest associations (**Figure 13C**). A ratio intensity plot revealed differences in mean hydrogen bond occupancies for each nucleotide-binding model (**Figure 13D**) with alterations in the mean number of hydrogen bonds tracked for each domain-domain interaction during the simulation time (**Figure 13E**). Two exceptions to the increased inter-domain bonding observed in the NLRP3ΔPYD-ADP trajectories were identified for the NBD to HD1 as well as the NBD to WHD. Small disparities were observed between models for the NBD-HD1 pairing, and the number of hydrogen bonds was comparatively increased for NBD-WHD during the NLRP3-ATP simulations. The highest number of internal bonds for both models occurred between the WHD and HD2, followed by bonds between the HD2 and LRR, with substantial decreases observed for the ATP-bound NLRP3 trajectories observed in both cases (**Figure 13E**). Concurrently, hydrogen bonding increased between the NBD and WHD in the ATP-bound model, as ATP binding resulted in an opening of the structure and increased the proximity of the NBD to the WHD. Several residues involved in hydrogen bonding between the WHD and HD2 in NLRP3ΔPYD-ADP are replaced by NBD-WHD interactions in NLRP3ΔPYD-ATP. Ultimately, fewer stabilising interactions exist between the WHD and HD2, with no bonds identified between the WHD and HD2 after 290 ns for the NLRP3ΔPYD-ATP simulation. The opening mechanism also results in considerable decreases in NLRP3ΔPYD-ATP hydrogen bonding between the HD1 and LRR, as well as HD1 and HD2. Furthermore, bonds between the NBD and HD2 were lost completely following the equilibration of the ATP-bound complex. Taken together, the data support the importance of interactions between the ATP-bound NBD and the WHD in driving the open conformation observed for active NLRP3.

**Figure 13.**
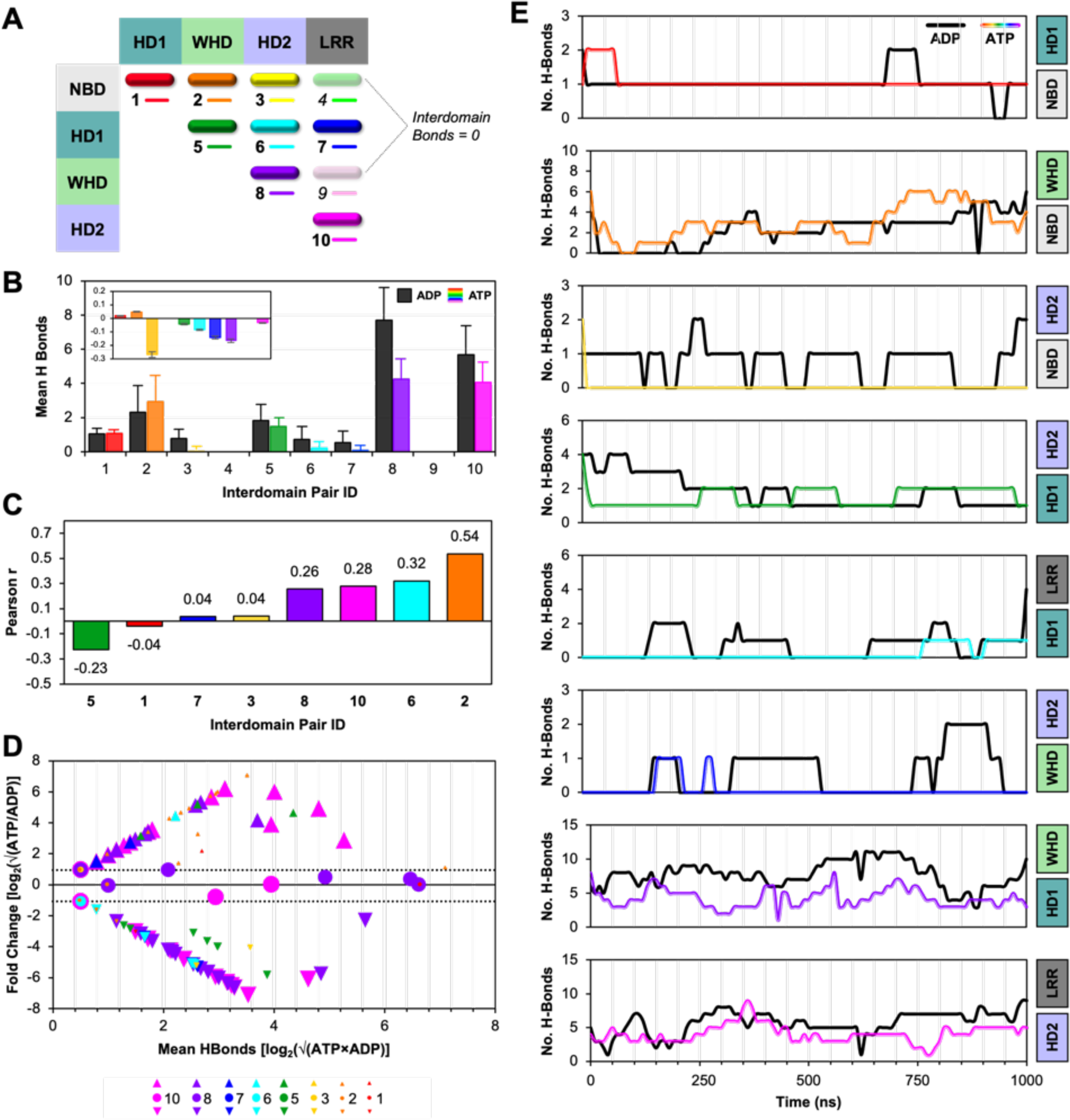
Internal Subdomain Hydrogen Bonding Within NLRP31′PYD. (**A**) NLRP3 domains, including the Nucleotide-binding Domain (NBD), Helical Domain 1 (HD1), Winged Helix Domain (WHD), Helical Domain 2 (HD2) and the Leucine Rich Repeats (LRR), are shown in a matrix with the associated colour and assigned interdomain bond pair ID number utilised in subsequent plots and bonds in the graphical representations of NLRP31′PYD, where colour-coding for bonds between specific subdomains are conserved in later panels. (**B**) The mean ± SD number of hydrogen bonds between domains are listed by region interaction ID, with ADP bars in black and ATP bars in the colours shown in panel (A). Inset; ATP/ADP fold change values shown for each interdomain pair, y-axis runs from -0.3 to 0.2. (**C**) Pearson correlation coefficients (r) were calculated for the number of bonds formed between domains as a function of the simulation time, shown here as a bar plot with data labels sorted by value from negative to positive. Values close to 1 indicate perfect correlation, while negative values represent negative correlation between ADP- and ATP-bound models. (**D**) Ratio intensity plot of the mean number of hydrogen bonds observed in the ADP- and ATP-bound NLRP31′PYD simulations, highlighting the differences between all internal hydrogen bonds in both systems. Data were transformed onto the y axis by the base 2 logarithm of the occupancy of the bond during the NLRP31′PYD-ATP trajectory over the occupancy of that same bond in the ADP trajectory, and onto a mean average scale (x-axis) with the base 2 log of the square root of the product of both occupancies. Values above 1 indicate an increase in that hydrogen bond in the ATP-bound NLRP31′PYD simulation (up arrows), while values below -1 (down arrows) indicate the reverse. Symbols are sized to allow for visual distinction between interaction groups and not due to data values. In (**E**), the mean number of hydrogen bonds for each domain-domain interaction is shown as a function of simulation time from 0 to 1000 ns. All ADP data are plotted as black lines, while the ATP data are plotted in the interaction ID colour shown in panel A.

Several specific residues displayed changes in domain interactions with nucleotide occupancy (**Figure 14**). Specifically, Gln480 and Glu527 residues within the WHD provided stable bonds with the NBD over the time course of the NLRP31′PYD simulation with ADP, yet both were repositioned to bond with the HD2 in the ATP-bound NLRP31′PYD model. Two stable hydrogen bonds between the NBD and WHD were formed by Arg351 to Gln480 (**Figure 14A**) as well as from Arg351 to Glu527 (**Figure 14B**) during the final ∼800 ns of the NLRP3ΔPYD-ADP simulations. These, along with bonds from both Gln308 and Gly309 to Gln480, were lost when Gly527 and Gln480 were repositioned to form bonds with the respective HD2 residues Leu562 and Lys552 in the NLRP31′PYD-ATP simulation.

**Figure 14.**
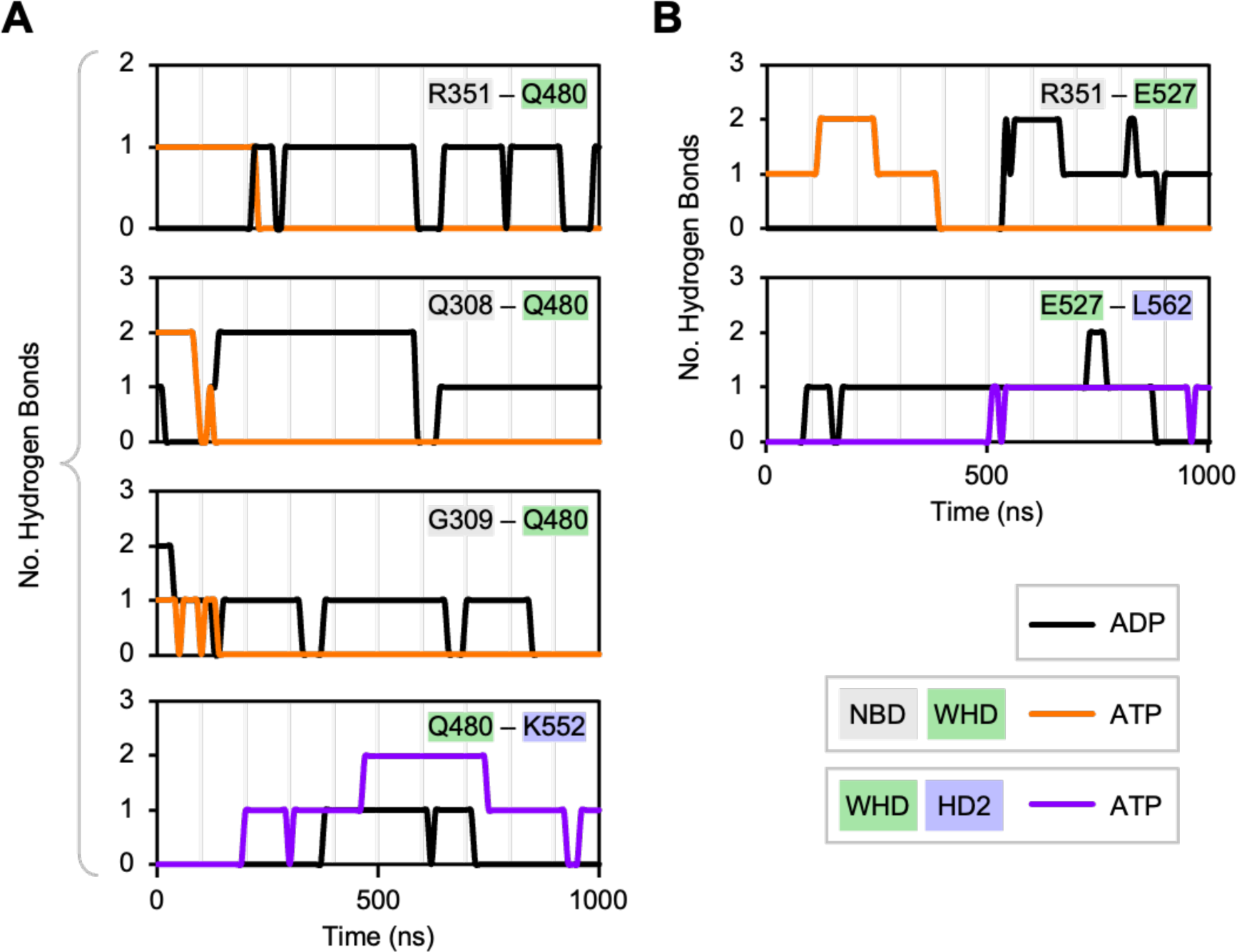
NBD-WHD Interdomain Hydrogen Bonds Are Replaced by HD2-WHD Bonds with ATP Occupancy. Interdomain hydrogen bonding within NLRP3 was observed for the nucleotide-binding domain (NBD), Winged Helix Domain (WHD), and Helical Domain 2 (HD2). In (**A**), Q480 (WHD) bonds to NBD residues R351, Q308 and Q309 are lost during the simulation time as Q480 binds K552 (HD2). In (**B**), the E527 (WHD) bond to R351 (NBD) is lost when E527 repositions to bind L562 (HD2). Legend indicates domain and colours where labels are highlighted according to residue domain and lines are coloured in black (ADP) or orange/purple according to the matrix shown in Figure 13.

### Structural Overview of Collective Motions for NLRP3-ΔPYD with Nucleotide Binding

Essential dynamics analyses provide a means to visualise and parse the collective motions of molecular simulations over time. In **Video 2** (http://dx.doi.org/10.17632/f5tf9rcxgc.1) a comprehensive view of the first three dynamic projections is shown individually and coloured by domain and subdomain. Six projections (i.e., PC1-PC3 for both NLRP3ΔPYD-ADP and NLRP3ΔPYD-ATP; **Figure 15A** and **15B**) are shown together at the conclusion of the video to highlight the considerable distinctions in dynamics driven exclusively by bound nucleotide identity. In **Figure 15C-E**, a comparative view of the collective motions for principal components (PC1-PC3) are represented by vectors from all the Cα backbone atoms and the centre of mass (COM) of each domain and subdomain. The collective motions presented in **Video 2** and **Figure 15** demonstrate global rearrangements highly dependent on the N-terminal PYD-NACHT linker and the acidic loop (AL) region within the LRR.

**Figure 15.**
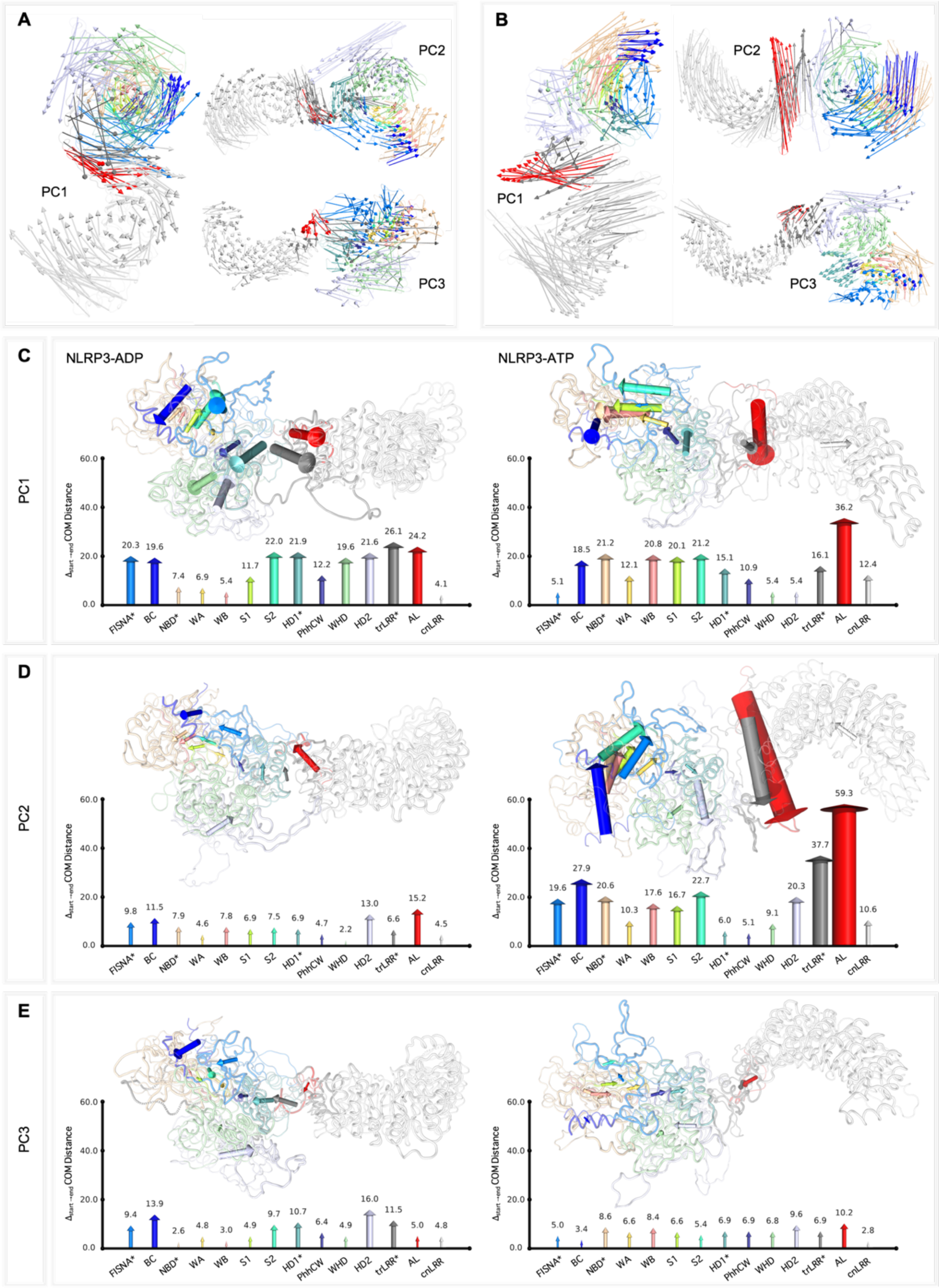
Structural Representation of NLRP31′PYD-ADP/ATP Essential Dynamics. Vectors representing the collective motions of each Cα atom along the first three eigenvectors (ADP-bound, **A**; ATP-bound, **B**) are shown coloured by domain as indicated in subsequent panels. Starting structures are represented as highly transparent ribbon structures to emphasise the vectors. Vectors are drawn from the centre of mass (COM) of each domain and subdomain at the starting position to the ending position. Arrows’ widths and heights were adjusted against all other distances to provide a comparative view of the collective motions of the major principal components (PC1, **C**; PC2, **D**; PC3, **E**). Labelled arrow charts indicate the domain or subdomain for which the vectors were calculated. Note: the FISNA (Fish-Specific NACHT-associated domain), NBD (Nucleotide-binding Domain), HD1 (Helical Domain 1) and trLRR (transition LRR) subdomains were not included in the COM or distance calculations and can instead be viewed individually as arrows drawn for the Basic Cluster (BC); Walker A, B (WA/WB) and Sensor 1 (S1) motifs; and Acidic Loop (AL), respectively. A script obtained from the open-source Pymol repository [modevectors.py] was modified to generate the vector representations.

Shorter 100 ns MD production runs of WA, WB, P412A and R262W mutants highlight similar motions and conformational changes observed for inactive, wild-type ADP-bound NLRP3 and the inhibitory WA, WB and P412A mutants. Intriguingly the R262W mutant underwent similar opening mechanisms to those of ATP-bound NLRP31′PYD (**Video 5;** (http://dx.doi.org/10.17632/f5tf9rcxgc.1). Furthermore, the closing/opening mechanisms were largely unaffected by bound nucleotide identity, confirming the importance of these regions and residues in regulating the global conformation of NLRP3.

## 4. DISCUSSION

### In this study, MD simulations of NLRP3 protein topology were completed as a function of bound nucleotide

**Simulations of** NLRP31′PYD with ADP indicated relatively stable conformations and few global rearrangements (i.e., as shown by decreased protein RMSD, Rg, SASA, and solvent accessibility) during the MD time-course. The ADP-bound cryo-EM structural template (i.e., 6NPY (Sharif et al. 2019)) on which these models were built was solved in the presence of extensive NEK7 interactions. Interestingly, the removal of these constraints over the course of the microsecond long simulations resulted in relatively stable trajectories and minimal large-scale domain reorganizations for the ADP-bound NLRP31′PYD model. The binding of ATP was found to induce marked structural transitions in the NLRP3 protein backbone that were distinct from those observed with ADP binding. Indeed, the computed mean RMSF values were increased for every domain and subdomain of NLRP31′PYD with ATP occupancy as compared with ADP. Validation of the concatenated trajectories provided a high degree of confidence in the sampling convergence achieved in these simulations, as measured by the converging RMSD values, stable and concordant nucleotide trajectories relative to the protein, favourable overall MM-PBSA ligand binding energies and low cosine coefficients of the principal eigenvectors obtained from ED analysis.

The simulations also suggest distinct binding modes for ADP and ATP within the NBD of the NLRP31′PYD model (**Figure 16**). The β-phosphate moiety of ADP protruded outward from the core cavity of NLRP3, with the adenine moiety deeply buried inside the catalytic cleft. This nucleotide orientation was similar to that previously observed for NOD2 in computational docking studies (Maharana et al. 2015) and in the ADP-bound crystal structure (Maekawa et al. 2016). In contrast, ATP showed a reverse binding mode aligned with the orientation previously reported for NLRC4 (Hu et al. 2013), with the adenine moiety protruding outward and the γ-phosphate group toward the core cavity. These nucleotide orientations can be observed more clearly for NLRP3 in **Video 4**, where x and y-rotated views of the binding pocket for the top ADP- and ATP-bound clusters are shown.

**Figure 16.**
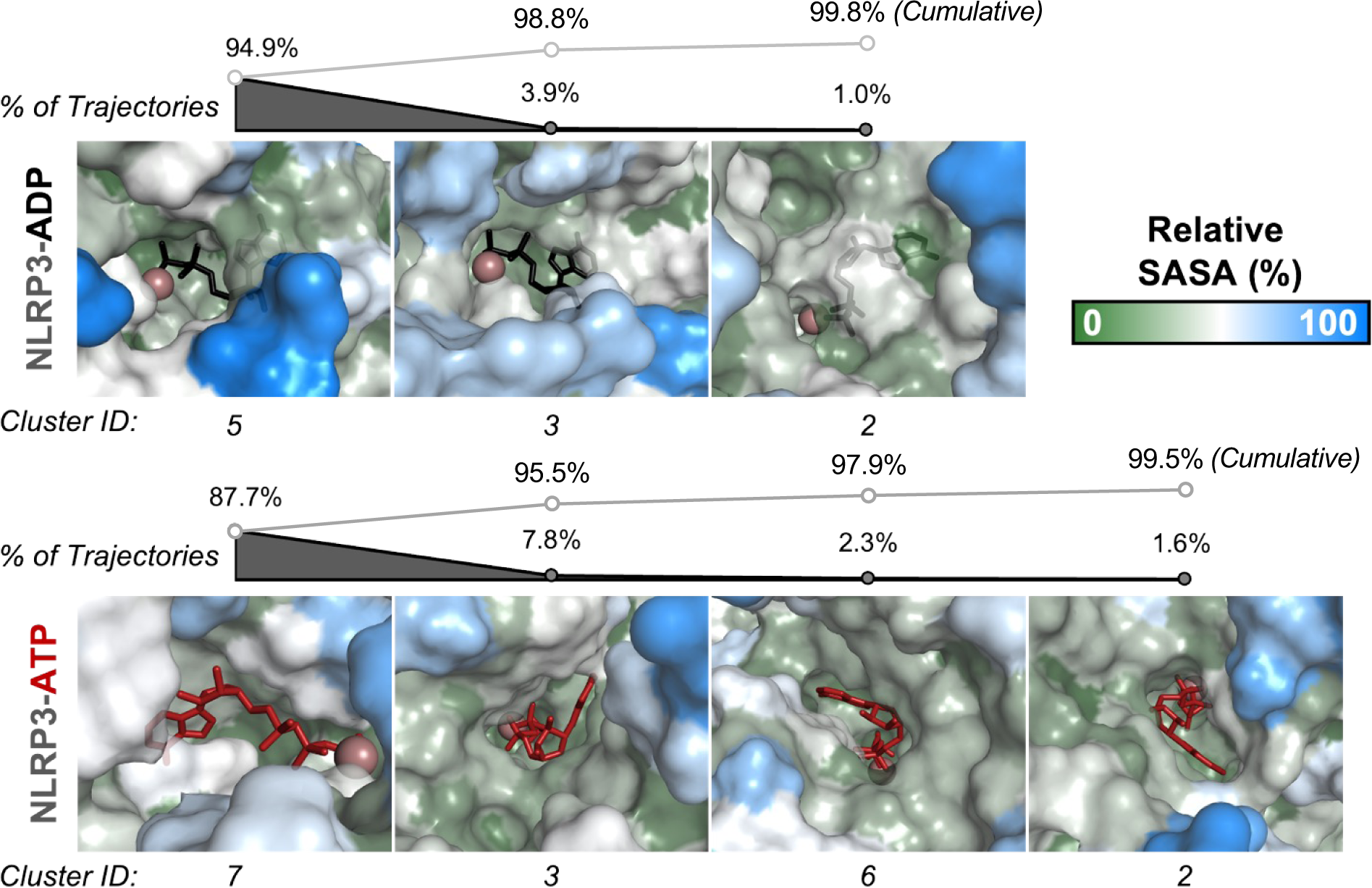
Binding Modes of ADP and ATP in NLRP31′PYD. Representative views of the nucleotide binding pockets of the top cluster medoids identified for NLRP31′PYD-ADP (top) and NLRP31′PYD-ATP (bottom). The percentage of the trajectory represented by each medoid is represented above each view, and cumulative percentages are listed above the grey lines. Surface views of the NLRP3 protein are coloured by relative solvent accessible surface area calculated per residue and shown in a spectrum from blue to white to green representing 0 to 100% solvent accessibility. ADP and ATP are coloured black and dark red, respectively. Mg^2+^ is represented as a violet sphere.

Ultimately, the MD simulation data support a mechanism for NLRP3 activation that involves: (1) a relaxation of the NBD core architecture upon ATP binding; (2) successive molecular rearrangements that occur after ATP occupancy to increase the proximity of the NBD to the WHD region; (3) subsequent increases in intramolecular NBD-WHD contacts with concomitant weakening of interactions between the WHD and both the HD1 and HD2 regions; (4) release of WHD to enable hinging of the HD2 and LRR domain; and finally, (5) migration to an open NACHT conformation that is associated with the activated form of NLRP3. These conclusions are concordant with recent structures of multimeric NLRP3, which require specific positioning between the NACHT and LRR domains for nucleation of the inflammasome complex (Hochheiser et al. 2022; Andreeva et al. 2021; Ohto et al. 2022). Moreover, these structures of inactive multimers suggest that the PYD domain of NLRP3 is buried in the central pore of the homo-oligomeric ring prior to ATP binding. So, the mechanism described above and highlighted in **Figure 17** provides a basis for the opening of the NACHT and rearrangement of the NACHT-PYD linker region to expose the interface required for homotypic ASC recruitment and NLRP3 inflammasome activation following ATP binding.

**Figure 17.**
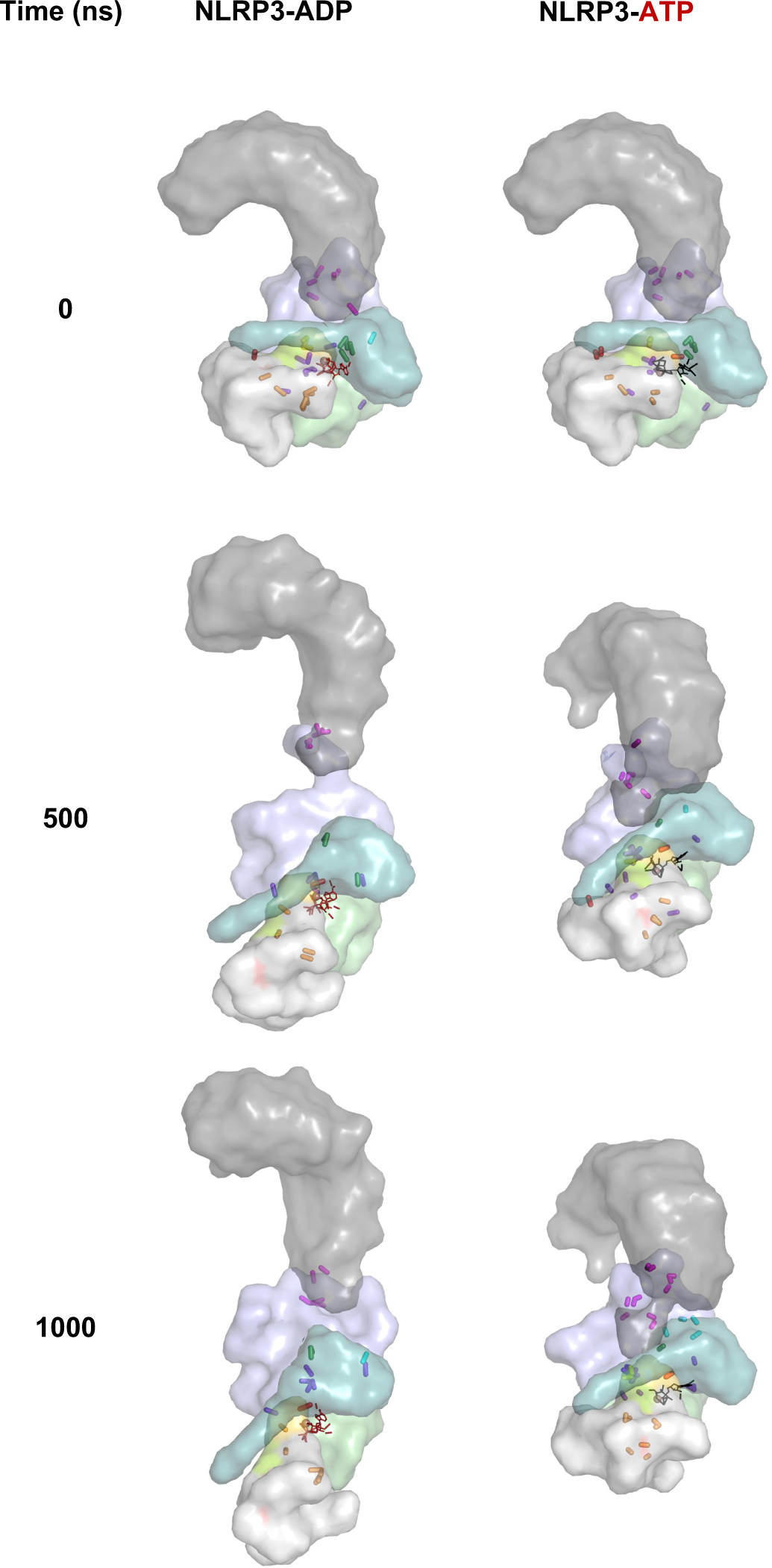
Nucleotide-occupancy Drives Global Conformational Changes in NLRP3. Transparent surface representations of NLRP31′PYD-ADP (sticks in black) and NLRP31′PYD-ATP (sticks in dark red) are shown at start (i.e., 0 ns), 500 ns and 1000 ns. Domains and bonds are coloured according to the interdomain matrix supplied in Figure 13. Images are snapshots sampled from Video 3.

Both HD1 and HD2 regions of NLRP31′PYD produced substantive molecular motions during the MD simulations with a prominent hinging action observed for the HD2 backbone. The molecular motions resulted in similar displacement distances and angles between NACHT and LRR domains to those defined in the activated NLRC4 structure. The hinging between the HD2 and LRR was previously postulated to be the driving mechanism by which the NACHT domain adopts an open conformation during NLRP3 inflammasome activation (Tapia-Abellan et al. 2021). The presented data support this hypothesis and provide detailed accounts of the inter- and intra-molecular interactions that accompany the hinging motions. An interesting finding is that ATP binding alone can support an activation mechanism for NLRP3 under dynamic simulations with rather subtle deviation of critical residues within the NBD conveying substantive conformational changes on to the global structure.

NLRP3 oligomerization is considered central to inflammasome assembly and activation of downstream signaling events. Yet, recent cryo-EM findings also support the existence of inactive oligomeric NLRP3 complexes (PDB IDs: 7LFH (Andreeva et al. 2021), 7PZC (Hochheiser et al. 2022), 7VTP (Ohto et al. 2022) and 7VTQ (Ohto et al. 2022)). These published structures highlight several key topographic features that correlate well with the ADP-bound NLRP31′PYD simulation data. The dynamic motions minimised at energy basins in MD simulations of NLRP31′PYD with ADP are reflective of similar loop conformations identified within the homodecameric structures (Sandall, Ziehr, and MacDonald 2020). Moreover, these data provide evidence that ADP-bound and inactive NLRP3 may require minimal supporting cofactors under basal conditions to initiate the formation of thermodynamically stable inflammasome complexes. A recent supportive publication indicates ADP-bound NLRP3 can form homo-oligomeric decamers in the presence of a small inhibitory molecule, MCC950, suggesting this conformation occurs at the basal level prior to effector recruitment and activation (Hochheiser et al. 2022). As inactive NLRP3 was shown to oligomerize in multiple studies (Hochheiser et al. 2022; Andreeva et al. 2021; Ohto et al. 2022) (37, 164,188), this suggests that both ADP and ATP have important yet, distinct, impacts on NLRP3 conformations within the multimeric state. These prior structural analyses concluded that the PYD domains of the NLRP3 subunits were buried within the centre of the decamer ring, and no effector molecules were associated with this subdomain structure. High atomic fluctuations in the FISNA domain identified during MD simulations with either ADP or ATP suggest this region contributed to collective motions that concentrated on the NACHT. The FISNA region was previously hypothesised to drive an opening of the homodecamer to provide exposure of internally buried PYD domains within the ring (Hochheiser et al. 2022). Highly divergent dynamics observed within this region during MD simulations of ADP- and ATP-bound NLRP31′PYD end further support for this mechanism of transformation.

Theoretical calculations of free energies by MD are valuable for assigning variable conformational states that are undetected by empirical experimentation (Christ, Mark, and van Gunsteren 2010). Alchemical free energy (AFE) methods are the gold standard in theoretical free energy calculations; however, these Monte Carlo-based methods require abundant sampling of complexes, ligands, and intermediate states and are thus costly to compute. Convergence is especially challenging for systems such as NLRP3, where large structural reorganisations occur following nucleotide exchange (Meng, Dashti, and Roitberg 2011). Molecular mechanics energies combined with the Poisson–Boltzmann or generalised Born and surface area continuum solvation (MMPBSA and MMGBSA) methods achieve a better balance between computational efficiency and accuracy, have successfully reproduced experimental findings, and are employed frequently in analyses of ligand-protein complex MD simulations as well as in ligand docking (Wang et al. 2019). In this study, the Poisson-Boltzmann (PB) method employed was non-trivial in terms of computational cost due to the large molecular size of NLRP3, but effective simulations were achievable with available resources. MM-PBSA outputs defined the critical residues in ADP and ATP binding and support a complex integration of stabilising interactions to increase the favourable electrostatic energies of NLRP31′PYD with ATP binding.

The collective motions presented in **Figures 8** and **15** demonstrate global rearrangements that also involve the acidic loop (AL) within the LRR domain. So, this molecular space may provide additional opportunity for NLRP3 inhibition. Targeting the cleft provided by the AL and the first transitional LRR (trLRR) in the inactive state to inhibit the opening and hinging mechanisms observed in the dynamic simulations of NLRP3. This region encompasses the α23-24 helices of the NLRP3 structure (labeled in **Figure 18**) and the small pockets created by the first leucine repeat of the LRR. These residues underwent relatively low fluctuations upon activation and present a stable target which retains a significant impact on the NACHT domain dynamics through interactions of the AL upon activation. Supposing critical hydrogen bonding patterns from the trLRR to the HD2 could be stabilised by a small molecule inhibitor within the approximate NLRP3 region covering residues 695 to 735, and the opening mechanism governed by fewer direct interactions between the HD2 and the LRR could be abrogated. Furthermore, this loop is known to play a role in coordinating multimerisation, as shown through the now available oligomeric structures of NLRP3 (Hochheiser et al. 2022; Andreeva et al. 2021; Ohto et al. 2022). Small molecules designed to target this region could be computationally optimised to maximise the probability of a specific and potent antagonism, and additional MD simulations of potential inhibitors would provide a pipeline to advance drug discovery endeavours.

**Figure 18.**
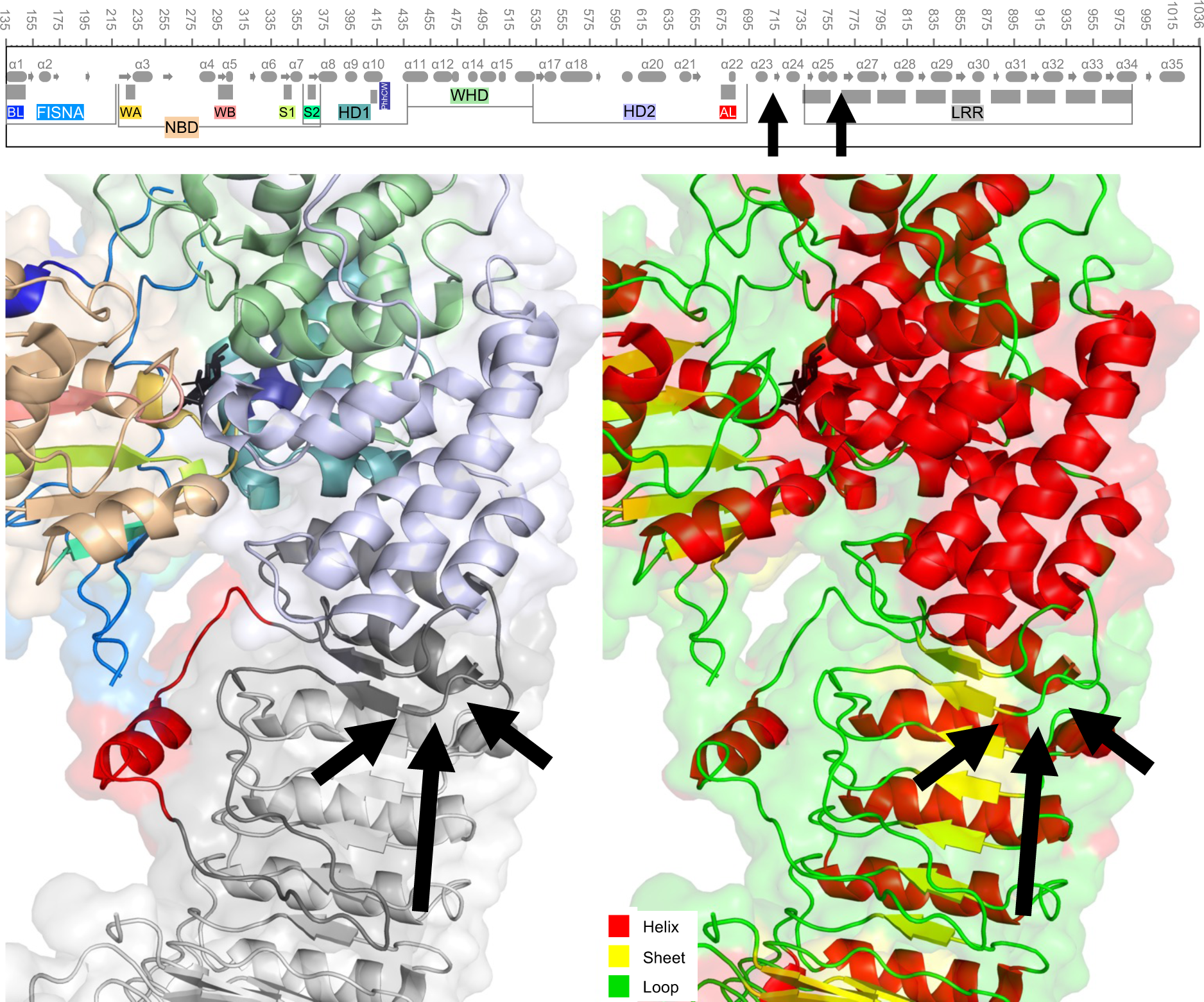
Novel Locational Target for Intervention of NLRP3 Activation. Cartoon representations of ADP-bound NLRP31′PYD are shown coloured by domain and motifs (left) and by secondary structure (right). Black arrows point to the region on the structures between the α23-24 helices of the NLRP3 structure (residues 695 to 735), and on the secondary structure diagram above the protein structures. Domains are coloured as in the labels of diagram, while secondary structure elements are coloured in red for helices, yellow for sheets and green for unstructured loops.

## ACKNOWLEDGEMENTS

This work was supported by a grant from the Natural Sciences and Engineering Research Council of Canada (NSERC; RGPIN/04379-2019 to JAM). CFS was also holder of an NSERC PGS-B Graduate Scholarship Award.

## SUPPLEMENTARY ONLINE MATERIALS

Several videos are referenced in this work; available for download from the Mendeley Data Repository at the following web address: http://dx.doi.org/10.17632/f5tf9rcxgc.1

**Video 1. MD trajectories displaying the NLRP31′PYD-ADP and NLRP31′PYD-ATP complexes during a 1 μs molecular dynamics simulation.** Nucleotides are shown as sticks, coloured black (ADP, left) or red (ATP, middle). Mg^2+^ ions are shown as spheres in violet. Domains are coloured as shown in the legend. A merge view is shown to the right.

**Video 2. NLRP3 Essential Dynamics: PC1-PC3 projections of NLRP31′PYD-ADP and NLRP31′PYD-ATP over two separate 1 μs simulations.** Structural projections of PC1, PC2 and PC3 throughout the MD simulations for ADP and ATP bound NLRP31′PYD conformational dynamics, shown as backbone Cα ribbon diagrams. For each PC, the starting structure is displayed, then vectors from the starting Cα atoms to the end position are added, coloured according to domain. The center of mass (COM) of NLRP3 domains and motifs are displayed as spheres, sized according to the overall translation distance from COM_start_ to COM_end_, and subsequently the COM_start➝end_ are shown. Arrow charts with labeled distances in angstroms are shown concurrently. Finally, the structural projection is shown with 50 representative frames, first with COM vectors and then without. Videos cycle from start to end and back to start. Additionally, PC1-3 projections for both nucleotide-bound structures are shown together in the final 22 seconds of the video (3:20). A script obtained from the opensource Pymol repository [modevectors.py] was modified to generate the vector representations.

**Video 3. Subdomain bonds in the NLRP31′PYD-ADP and NLRP31′PYD-ATP structures during a 1 μs molecular dynamics simulation.** Surface representations of the ADP-(left) and ATP-(right) bound NLRP31′PYD structures are coloured by domain as shown in the legend. The WA (gold), WB (pink) and S1 (green) motifs are coloured within the grey NBD. Interdomain bonds are shown as thick lines and coloured as shown in the matrix on the bottom left. Direct hydrogen bonds are shown as thin lines and coloured black (ADP) or red (ATP). The Mg_2+_ ion is shown as a sphere in violet.

**Video 4. Nucleotide SASA of Top MD Clusters.** Surface representations of the ADP (top) and ATP (bottom) nucleotide binding sites are oriented to demonstrate the solvent accessibility of the nucleotide binding sites. Residues are coloured by relative solvent accessible surface area from dark green (0%) to white (50%) to blue (100%) as indicated. Then, views zoom to 4 Å away from the center of mass of the nucleotide, and 360° rotations around the y-axis, and then x-axis, are shown. In the smaller video panels, the rotation of the entire structure is shown for reference. Surface views for residues greater than 40 Å from the nucleotide are hidden and shown as transparent backbone ribbons for clarity.

**Video 5. MD trajectories of the WT-NLRP31′PYD protein alongside the WA, WB, P412A and R262W mutants.** Left: ADP bound structures, right, ATP bound structures. Top panel, cartoon and stick representations of the nucleotide binding pocket, where ADP is shown in black and ATP is shown in red. The Mg^2+^ ion is shown as a sphere in violet. Mutated residues are shown coloured by domain in the WT panel, with mutant residues shown in magenta in the subsequent panels. Mutant simulations were run for 100 ns each, with WT videos trimmed to the same. Videos repeat once.

